# Autoimmunity to hypocretin and molecular mimicry to flu antigens in Type 1 narcolepsy

**DOI:** 10.1101/378109

**Authors:** Guo Luo, Aditya Ambati, Ling Lin, Mélodie Bonvalet, Markku Partinen, Xuhuai Ji, Holden Terry Maecker, Emmanuel Jean-Marie Mignot

**Affiliations:** Center for Sleep Sciences and Medicine, Stanford University School of Medicine, Palo Alto CA 94304, USA; Helsinki Sleep Clinic, Vitalmed Research Centre, Helsinki, Finland and Department of Clinical Neurosciences, University of Helsinki, Helsinki, Finland; Immune Monitoring Center, Institute for Immunity, Transplantation, and Infection, Stanford University School of Medicine, Palo Alto, CA 94305, USA

## Abstract

Type 1 narcolepsy (T1N) is caused by hypocretin (HCRT) neuronal loss. Association with the Human Leukocyte Antigen (HLA)-DQB1*06:02/DQA1*01:02 (98% vs 25%) heterodimer (DQ0602), T cell receptor (TCR) and other immune loci suggest autoimmunity but autoantigen(s) are unknown. Onset is seasonal and associated with influenza A, notably pandemic 2009 H1N1 (pH1N1). An extensive unbiased DQ0602 binding peptide screen was performed encompassing peptides derived from Pandemrix^®^ X-179-A pH1N1 influenza-A vaccine, a known T1N trigger, other H1N1 strains, and potential human autoantigens HCRT and RFX4, identifying 109 binders. The presence of cognate tetramer-peptide specific CD4^+^ T cells was studied in 35 narcolepsy cases and 22 DQ0602 controls after expansion of antigen-specific cells in Peripheral Blood Monocytes Cell (PBMC) cultures. Higher reactivity to influenza epitopes pHA_273-287_ (pH1N1 specific) and PR8 (H1N1 pre 2009)-specific NP_17-31_ were observed in T1N. Extensive reactivity to C-amidated but not native version of HCRT_54-66_ and HCRT_86-97_, which are two highly homologous peptides (HCRT_NH2_) was observed with higher frequencies of specific T cells in T1N. TCRα/β CDR3 sequences found in pHA_273-287,_ NP_17-31_ and HCRT_NH2_ tetramer positive CD4^+^ cells were also retrieved in single INFγ-secreting CD4^+^ sorted cells stimulated with Pandemrix^®^, confirming immunodominance and functional significance in DQ0602-mediated responses and molecular mimicry. TCRα/β CDR3 motifs of HCRT_54-66_ and HCRT_86-97_ tetramers were extensively shared. Particularly notable was sharing across subjects of an CDR3α, CAVETDSWGKLQF (in association with various CDR3β that used TRAJ24, a chain modulated by Single Nucleotide Polymorphism (SNPs) rs1154155 and rs1483979 associated with T1N. Sharing of CDR3β CASSQETQGRNYGYTF (in association with various CDR3α was also observed with HCRT_NH2_ and pHA_273-287_-tetramers across subjects. This segment uses TRBV4-2, a segment modulated by narcolepsy-associated SNP rs1008599. Higher HCRT_NH2_ positive CD4^+^ T cell numbers in T1N together with sharing of J24 CAVETDSWGKLQF in HCRT_NH2_ autoimmune responses, indicates causal DQ0602-mediated CD4^+^ autoreactivity to HCRT in T1N. Our results provide evidence for autoimmunity and molecular mimicry with flu antigens modulated by genetic components in the pathophysiology of T1N.

## Main text

Whereas genetic^1–6^, epidemiological^7–11^ and pathophysiological^12–14^ studies implicate autoimmunity in response to flu infection in type 1 narcolepsy (T1N), a disease caused by hypocretin (HCRT) neuronal loss^15–17^, the identity of the autoantigen is unknown. Potential autoantigens explored have included HCRT sequences^18–21^, HCRT receptor 2 sequences^22^ and the protein TRIB2^23^ but these results have either been withdrawn^24^ or not replicated^25–27^. Similarly, staining of hypothalamic slices with T1N sera has been found negative^28^.

Leading on from prior work^1–6^ a large recent GWAS analysis of over 4,500 patients across multiple ethnic groups confirmed primordial importance of the Human Leukocyte Antigen (HLA) DQ0602 heterodimer in disease predisposition, with important secondary associations in T cell receptor (TCR) loci α and β and other immune loci such as perforin. This suggests importance of DQ0602 presentation of antigens to CD4^+^ by Antigen Presenting Cells (APC) and likely subsequent CD8^+^ destruction of hypocretin cells^29^. Hypocretin cell cytotoxicity through CD4^+^ and CD8^+^ mediation has also been shown to cause narcolepsy in an animal model^30^.

Recently, much has recently been learned regarding what may trigger T1N autoimmunity. Onset of T1N in children, which can often be timed to the week, is seasonal and peaks in spring and summer^10^. Onset has also been associated with Streptococcus Pyogenes infections^9,31^ and Influenza-A^10^, suggesting the trigger may be a winter infection, followed by a delay peaking at 5 months once sufficient hypocretin cell loss has occurred and symptoms appear. Most strikingly, prevalence of T1N increased severalfold in China with a similar delay following the 2009-2010 “Swine Flu” H1N1 influenza pandemic (pH1N1)^10,32,33^, however, association with the pandemic is less clear in other countries^11,34^. Similarly, cases of T1N have been triggered by Pandemrix^®^ in Europe with relative risk increasing 5 to 14-fold in children and adolescents and 2 to 7-fold in adults after vaccination^33^. As Pandemrix^®^ is an AS03 adjuvanted vaccine containing the artificially produced reassortant strain X-179A, a mix of PR8, an old H1N1 strain derived from pre-2009 seasonal H1N1 but containing key H1N1 2009 surface proteins (Hemagglutinin-HA and Neuraminidase-NA)^35^, flu proteins are likely to be critically involved. The fact that HLA and TCR genetic associations are universal^6,36-40^ is also consistent with a flu trigger, as Influenza-A infections occur on gobal basis^41^. Importantly, however, even with Pandemrix^®^ vaccination in Europe, only 1 in 16,000 vaccinated children developed T1N. As has been illustrated above, both the H1N12009 pandemic and the Pandemrix^®^ vaccination have exibited variable effects across different countries, thus demanding the consideration of additional factors to fully explain T1N occurences.

## Results

### DQ0602 restricted epitopes of H1N1 flu responses in T1N versus controls suggest a role for pHA_273-287_ and NP_17-31_

Complementing our previously published data^18,24^, we screened overlapping 15-mers for DQ0602 binding for key proteins contained in X-179A and pH1N1 wild type, HA, NA, Influenza RNA-dependent RNA polymerase subunit 1 (PB1) and nucleoprotein (NP) (Supplementary Table 1). HA and NA were selected as these are key surface proteins enriched by design in flu vaccines such as Pandemrix^®^ (to facilitate B cell responses)^35,42^. PB1 was selected because it is of H1N12009 origin in Pandemrix^®^ and thus is also shared by the wild type H1N12009 virus and Pandemrix^®^, two known triggers of T1N^35,42^. We further added NP to our screen as it is the most abundant protein in flu and it is also present in large amounts in Pandemrix^®^ as enrichment process for surface proteins is not absolute in this vaccine (and many others, although some subunit specific vaccine were available, such as Focetria^®^ for pH1N1)^35^. NP of Pandemrix^®^ is of backbone PR8 origin^35,42^. Screening was primarily done using A/reassortant/NYMC X-179A (California/07/2009 x NYMC X-157, reference ADE2909) sequences complemented by wild type pH1N1 sequence (A/California/07/2009, reference AFM728) when polymorphism was evident, or sequences are of distinct origin (i.e. NP). Screening for these proteins identified large number of binders (Supplementary Table 1) using the biotin-conjugated EBV_486–500_ epitope (Bio-EBV, Bio-GGGRALLARSHVERTTDE), a known binder^18,43^. Peptides that outcompeted >75% of the reference EBV peptide signal were considered binders, and these were selected for tetramer screening^18,24^.

To assess for the presence of CD4^+^ T cells reactive to these epitopes, likely 9 mer-MHC class II binding repertoires were determined for all binders using three Immune Epitope Database prediction algorithms (SMM_align, NN_align, and consensus) and similarity scores. Peptides (15-mers) surrounding the cores and displacing most of the bio-EBV peptide in competition assays were used to construct tetramers by peptide exchange using biotinylated DQ0602 (Bio-DQ0602) cleaved with thrombin as described in the methods section. These tetramers were then used to FACS stain 10-day expanded peripheral blood mononuclear cells (PBMCs) cultures of 6 cases and 4 DQ0602 controls stimulated with Pandemrix^®^ (100 ng/ml) or the corresponding single peptide (6.25 μM) (see methods and staining results in Extended Data Figure 1a). The control and narcoleptic subjects were all known post Pandemrix^®^ vaccinees except for one case who developed T1N independent of vaccination 6 months prior to blood sampling (recent onset case).

Figure 1a shows the number of controls and T1N patients (from 10 subjects) positive for each tetramer staining as well as example staining (Figure 1d) with selected tetramers. Tetramer positivity ranged from no positive to 10 positives for some tetramers (Figure 1a), with mean number of positives per flu protein in post Pandemrix^®^ controls (4) and T1N (5) cases shown in Figure 1c. Post Pandemrix^®^ cases and controls were selected for Figure 1c comparison to ensure similar flu antigen exposure and as expected no difference was found across cases and controls for these antigens overall (all epitopes). In contrast, lower reactivity with PB1 versus HA, NP and NA was found, likely the consequence of the lower concentration of PB1 in the Pandemrix^®^ vaccine (and all vaccines) in comparison to these other proteins^35^.

**Figure 1.**
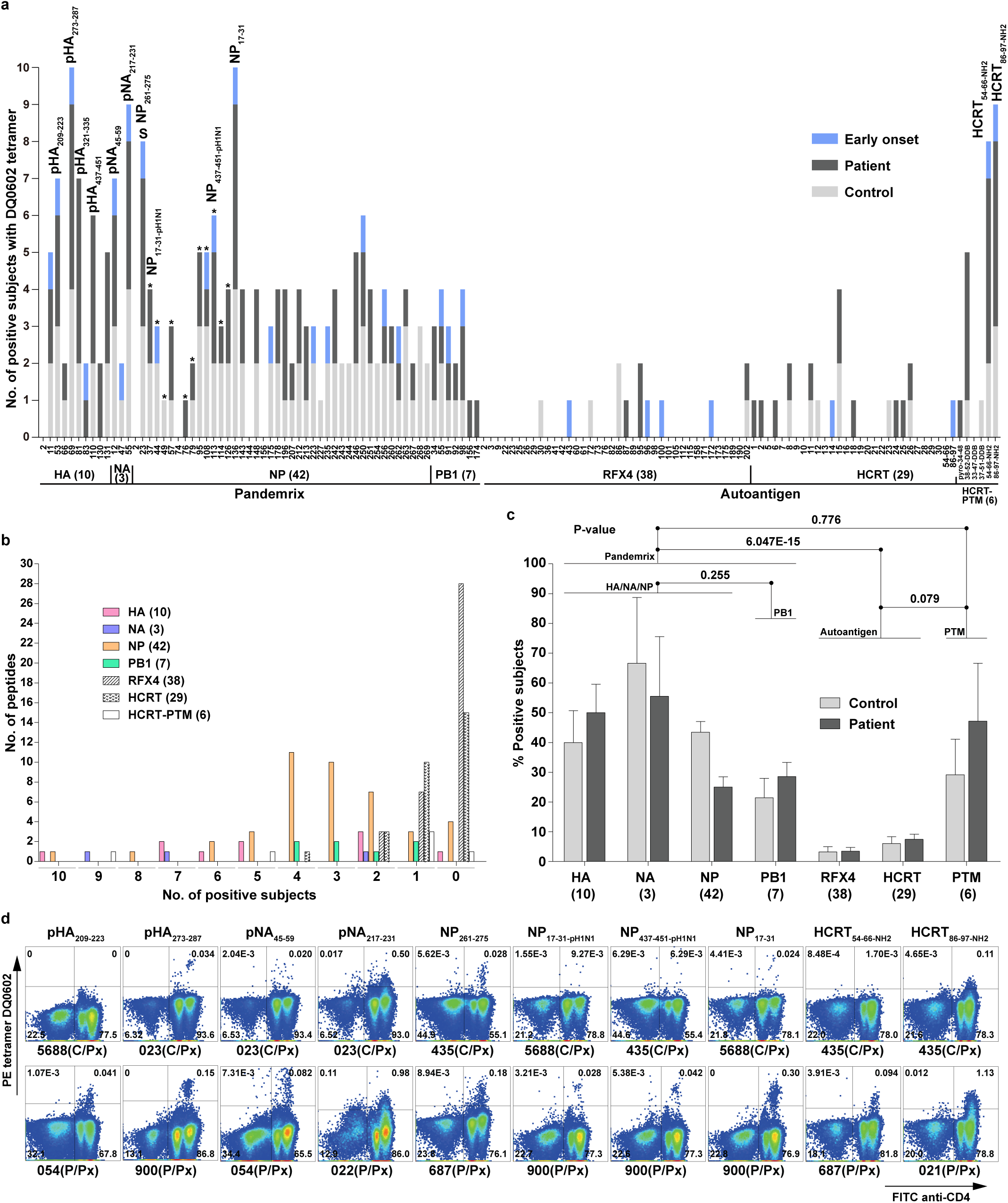
Results of primary tetramer screening in 10 subjects. **a**. Tetramer positivity in 5 post Pandemrix patients (P/Px), 4 post Pandemrix controls (C/Px) and one early onset patient (onset=0.3 year). Immunodominant epitopes are indicated. Number of peptides for each antigen is shown between brackets. PTM, post translational modification. S, shared by A/California/07/2009 and A/Puerto Rico/8/1934. * deriving from A/California/07/2009. For peptide and subject infomation, see Supplementary Table 1 and 9, respectively. **b**. Distribution of positive subjects per antigen. **c**. Mean percentage of subjects positive for DQ0602 tetramers in each antigen in patients and controls. Statistics between groups of antigens were calculated. **d**. Example of FACS staining obtained for various selected tetramers. CD3+ T cells are shown with frequency. The entire dataset of FACS staining plots is provided as Extended Data Figure 1. Note tail-like staining in many (but not all) cases with HCRT_86-97-NH2_ but not with other tetramers.

X179A derived tetramers with the most staining included pHA_273-287_, pNA_217-231_, NP_17-31_, NP_261-275_ (Figure 1d) suggesting these epitopes are immunodominant (Figure 1a, NP epitopes from pH1N1 instead of PR8 indicated by stars). Other immunodominant peptides include pHA_209-223_, pHA_321-335_, pHA_437-451_, pNA_45-59_, pNA_217-231_, NP_261-275s_ (shared epitope between pH1N1 and PR8), and NP_437-451_-pH1N1*. pHA_273-287_ is notable as it was previously reported to have homology with prepro-hypocretin sequences HCRT_54-66_ and HCRT_86-97_ and suggested be involved in a possible pathogenic mimicry using Enzyme-Linked ImmunoSpot (ELISpot) results, a publication that was retracted based on flawed ELISpot results (DQ0602 binding results were all verified)^18,24^. Interestingly, among immunodominant epitopes, responses to three epitopes pHA_273-287_ (of H1N12009 origin), NP_17-31_, and NP_261-275_ (of PR8 origin) were stronger in Pandemrix^®^ vaccinated patients versus controls and positive in the early onset cases (Figure 1a) and were thus selected for additional investigation.

Additional analysis of 24 cases and 14 DQ0602 positive controls confirmed significant overabundance of T cells positive for pHA_273-287_ and NP_17-31_ but not NP_261-275_ (Figure 2a and 2c, Extended Data Figure 1b and Extended Data Figure 2) suggesting that these two epitopes could be involved in the recruitment of differential CD4^+^ T cell populations related to T1N. Sub-analyses confirmed that these epitopes also gave significantly more response in a total of 11 post-Pandemrix^®^ cases (mean age 17±2yo) versus 8 post-Pandemrix^®^ controls (27±6 yo), an important control as these subjects had a more similar exposure to pH1N1 around the first wave of the pH1N1 pandemic infection (Extended Data Figure 2d). Findings with pNA_217-231,_ NP_261-275_ and NP_17-31-pH1N1_, the homologous sequence of PR8 NP_17-31_ in pH1N1 (2 amino acid difference in the core, see Supplementary Table 2) showed no differences overall and in subgroups (Extended Data Figure 2a-c). This demonstrates that increased flu reactivity in T1N versus controls is not broadly shared across all H1N1 epitopes, confirming screening results (Figure 1c).

**Figure 2.**
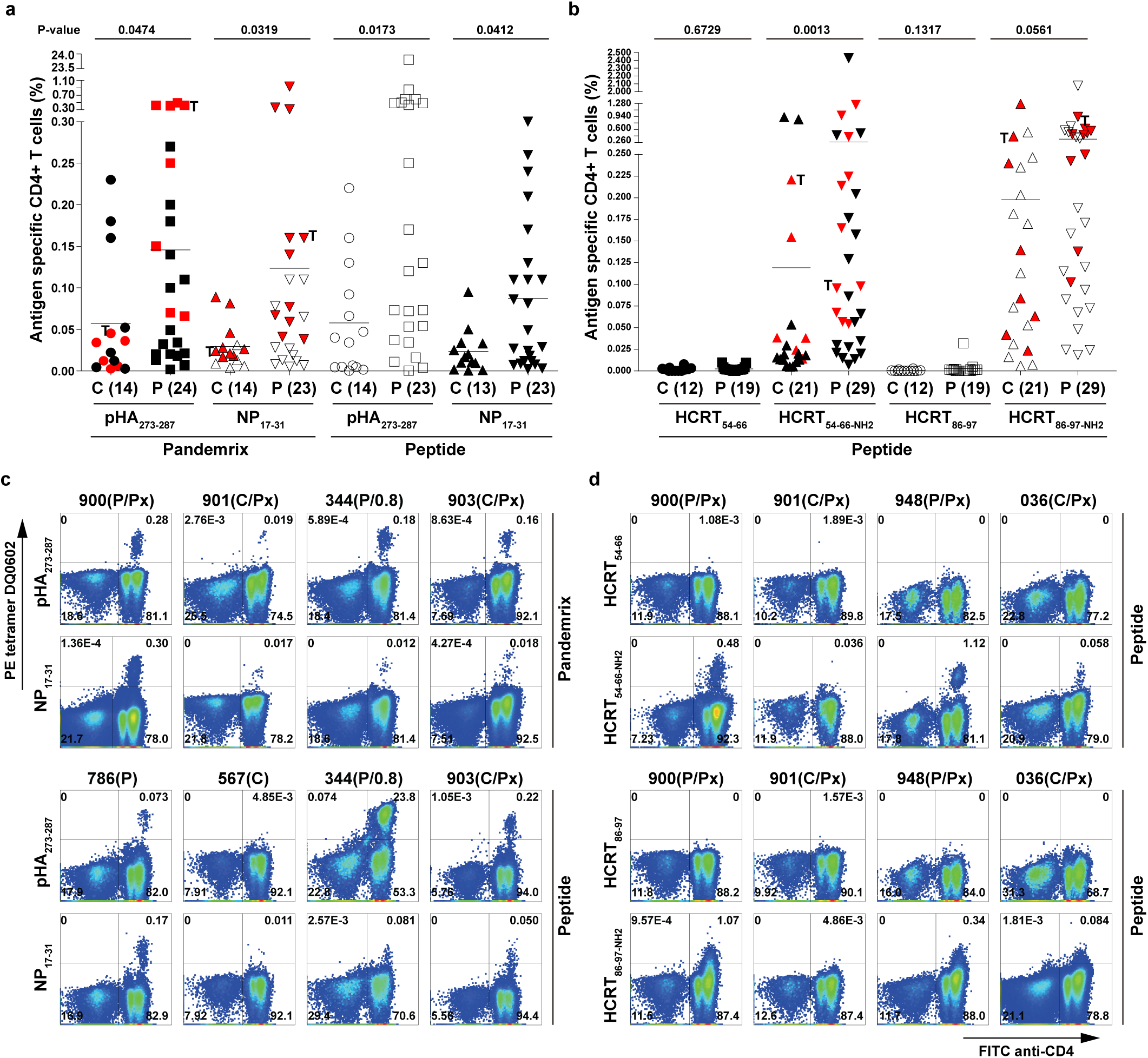
Results of secondary tetramer screening using pHA_273-287_, NP_17-31_, HCRT_54-66_, HCRT_54-66-NH2_, HCRT_86-97_, and HCRT_86-97-NH2_. **a**. Tetramer positivity of pHA_273-287_ and NP_17-31_ in controls versus patients stimulated with Pandemrix or peptide. **b**. Tetramer positivity of HCRT_54-66_/HCRT_54-66-NH2_ and HCRT_86-97_/HCRT_86-97-NH2_ in controls versus patients stimulated with peptide. **c**. Example of pHA_273-287_ and NP_17-31_ staining in a few subjects. **d**. Example of HCRT_54-66_, HCRT_54-66-NH2_, HCRT_86-97_, and HCRT_86-97-NH2_ tetramer staining in a few subjects (for all results, see Extended Data Figure 1). P, patient; P/Px, post Pandemrix patient; C, control; C/Px, post Pandemrix control. Interval is shown if patient is early onset (<1.0 year); T, twins; Red, single cell TCR sequencing. CD3+ T cells are shown in c and d with frequency.

### Broad CD4^+^ T cell autoreactivity to amidated HCRT_54-66-NH2_ and HCRT_86-97-NH2_ fragments when presented by DQ0602

We next tested whether T1N was associated with autoreactivity involving two proteins known to be primarily co-localized with hypocretin, hypocretin itself and RFX4, a transcription factor primarily expressed in testis and also exceptionally abundant in hypocretin cells^44^. Hypocretin and RFX4 binders were tested in the 10 subjects mentioned above, with results suggesting almost no or little cross reactivity, consistent with T cell tolerance toward autoantigen peptide sequences (Figure 1a-b). Autoreactivity was significantly lower than for all viral proteins (Figure 1c).

Because the pre-prohypocretin peptide is cleaved and post-translationally modified in the region where homologous sequences HCRT_54-66_ and HCRT_86-97_ are located (resulting in secreted hypocretin-1 and −2), most notably C-amidated, a transformation necessary to its biological activity, we also tested HCRT_54-66-NH2_ and HCRT_86-97-NH2_, modified peptides we have previously shown also bind DQ0602 with similar or higher affinity^18^. To our surprise, T cell reactivity to these fragments was high in both DQ0602 controls and T1N subjects (Figure 2b and Extended Data Figure 1). Of notable interest was the fact HCRT_54-66-NH2_ staining was generally restricted to distinct populations (not unlike what was observed with flu antigens) while HCRT_86-97-NH2_ staining was more heterogeneous, suggesting large amounts of low affinity tetramer cross-reactivity (Figure 2d and Extended Data Figure 1). These data were later expanded to 29 cases and 21 DQ0602 positive controls, and a significantly higher (p=0.0013) percentage of HCRT_54-66-NH2_ tetramer positive cells was found in T1N (Figure 2b), with marginally higher values for HCRT_86-97-_ NH2 (p=0.056). Sub-analysis of 13 post-Pandemrix^®^ cases (mean age 17±1yo) versus 10 post-Pandemrix^®^ controls (mean age 27±6 yo) confirmed significantly higher reactivity (p=0.0052) for HCRT_86-97-NH2_ and marginally higher values for HCRT_54-66-NH2_ (p=0.067) (Extended Data Figure 2f). Even more strikingly, in 7 recent onset cases (22±6 yo) versus 21 controls (23±3yo) higher reactivity was found to both HCRT_54-66-NH2_ and HCRT_86-97_ (non-amidated) suggesting an important role in the cause of T1N (Extended Data Figure 2g).

### T Cell receptor sequencing of flu tetramer positive cells indicate bias in TCRα/β usage consistent with *in-vitro* clonal expansion

Tetramer-reactive CD4^+^ T cells from 14-20 subjects were next FACS single sorted and sequenced in control and T1N patients (including the 11-13 post Pandemrix^®^ subjects and one early onset case mentioned above) for pHA_273-287_, NP_17-31_, HCRT_54-66-NH2_ and HCRT_86-97-NH2_. Sporadic additional sequencing (a few subjects, data not shown) was performed for other epitopes. Subjects sequenced are denoted in red in Figure 2a and 2b. For comparison, ∼10^6^ peripheral memory CD4^+^ T cells of the same subjects were sequenced for TCRα and TCRβ V and J segment baseline usage (data from: Ollila et al.^29^).

As expected, we found strong significant bias in TCRα/β usage (mostly for TRAV and TRBV segments) specific of each epitope, except for HCRT_54-66-NH2_ and HCRT_86-97-NH2_ where a large amount of sequence sharing was found across the two highly homologous HCRT epitope (also called HCRT_NH2_ as a group for future reference) and sequence usage diversity was higher in comparison to flu epitopes (Extended Data Figure 3, Supplementary Table 3). For example, TCRβ usage of pHA_273-287_ was notable for a most highly significant enrichment in TRAJ23 (Mantel Haenszel [MH] OR=13.6),TRAV13-1 (OR=5.4) and TRAV35 (OR=19.3) while TCRβ usage for the epitope was enriched in TRBV19 (OR=22.4) and TRBV4-3 (OR=35.3) in comparison to the same subject’s CD4+ cell peripheral usage, without differences in control versus T1N. Similarly, NP_17-31_ positive TCR cells were enriched in TRAV12-2 (OR=18.3), TRAV8-6 (OR=6.7), TRBV4-2 (OR=25.6), TRBV7-9 (OR=2.8), TRBV20-1(OR=4.7) (Extended Data Figure 3, Supplementary Table 3) without differences in control versus T1N. Of particular interest was enrichment in TRBV4-2, as T1N is strongly associated with rs1008599G (OR=0.81, p=6.63 10^−12^), a SNP that is the main eQTL for increased expression of this segment^29,45^.

### T Cell receptor sequencing of C-amidated HCRT tetramer positive cells indicate broad TCR reactivity and has consequently more limited bias in TCRα/β usage

Sequencing of amidated-HCRT positive tetramers gave a different picture (Extended Data Figure 3, Supplementary Table 3) in comparison to flu-positive tetramer. First, although increased usage of specific TCR segments was found, usage diversity was much higher (Extended Data Figure 3), notably when HCRT_86-97-NH2_ tetramers were used, giving a typically less distinct population with likely lower TCR affinity (Figure 2d, Extended Data Figure 1). In addition, in cases where HCRT_54-66-NH2_ and more rarely HCRT_86-97-NH2_ tetramers gave a clearly separated population on FACS plots (Extended Data Figure 1), sequencing resulted in almost pure oligoclonality of TCR sequences (even less diverse than with flu antigens) with minimal usage sequence sharing (patients 022, 948, 051 and 685 for HCRT_54-66-NH2_ and controls 567 and 391 for HCRT_86-97-NH2_ in Extended Data Figure 3, with corresponding FACS plots in Extended Data Figure 1).

### Shared CDR3 analysis and Grouping of Lymphocytes Interactions by Paratopes Hotspots (GLIPH) analysis across flu and HCRT_NH2_ tetramers

pHA_273-287_ and HCRT_54-66-NH2_/HCRT_86-97-NH2_ share weak epitope homology^18,24^, a result confirmed using http://tools.iedb.org/cluster/reference/^46^, with three Epstein Barr Virus (EBV) sequences as anchor (Figure 3e). Based on this, mimicry has been proposed but not demonstrated^18,24^. To search for mimicry across pHA_273-287_, NP_17-31_, HCRT_54-66-NH2_ and HCRT_86-97-NH2_, we first analyzed sharing of CDR3α, CDR3β and CDR3α/β pairs across epitopes. Supplementary Table 4 shows cross tabulations of shared sequences across epitopes.

**Figure 3.**
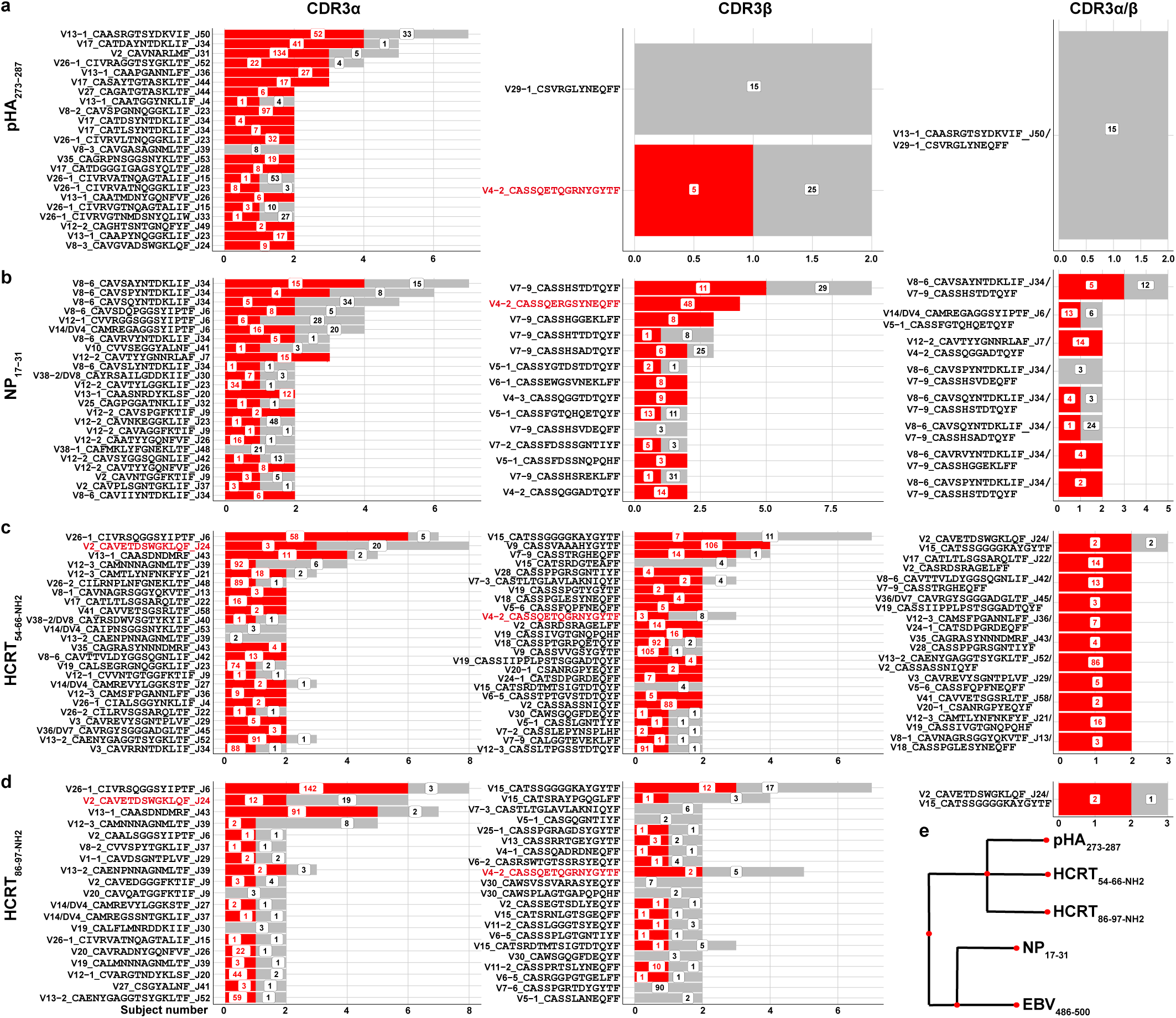
Shared CDR3α, CDR3β and CDR3α/β sequences for pHA_273-287_ (a), NP_17-31_ (b), HCRT_54-66-NH2_ (c) and HCRT_86-97-NH2_ (d) across subjects. Red, number of patients with corresponding CDR3 (per horizontal axis); Grey, number of controls with same CDR3 (per horizontal axis). Box numbers indicate total of times when the CDR3 occurred across patients and controls. **e**. pHA_273-287_, NP_17-31_, HCRT_54-66-NH2_ and HCRT_86-97-NH2_ sequence homology with EBV_486-500_ used as anchor. TRAJ24 and TRBV4-2 are labeled in red.

A large number of sequences were shared between HCRT_54-66-NH2_ and HCRT_86-97-NH2_ tetramers across CDR3α (11.8%), CDR3β (9.3%) and both CDR3 α/β (3.6%), a result that is not surprising considering that HCRT_54-66-NH2_ and HCRT_86-97-NH2_ only differ in their N-amidated C-terminal end, L-_NH2_ (hypocretin-1) and M-_NH2_ (hypocretin-2) in P9 of the binding repertoire (Supplementary Table 2). In contrast, pHA_273-287,_ a viral epitope which has been suggested to be a mimic of HCRT and has some homology with HCRT_54-66-NH2_ and HCRT_86-97-NH2_ (see Figure 3e) shared very few of the same clones with HCRT_54-66-NH2_ and HCRT_86-97-NH2_, for example only 1.9% and 1.5% for CDR3α, 0.7% and 1.8 % for CDR3β and none and 0.3% (2 instances) for CDR3α/β with HCRT_54-66-NH2_ and HCRT_86-97-NH2_ respectively. Potential sharing was even lower with NP_17-31_, which shared a few clones with pHA_273-287_, and at even lower level with HCRT_NH2_.

To explore differential CDR3 usage per epitopes and across narcolepsy versus control status we also used the Grouping of Lymphocytes Interactions by Paratopes Hotspots (GLIPH)_47_ in CDR3βs from pHA_273-287_, NP_17-31_ and HCRT_NH2_. A similar bioinformatic motif driven approach was also implemented for CDR3α. Tetramer derived unique CDR3s were compared to a large database of CDR3 sequences derived from bulk sequencing of ∼10^6^ CD4^+^ CD45RA-T cells obtained from the same 35 narcolepsy and 22 DQ0602 control subjects (see Supplementary Table 5 for a list of paratopes significant at the p<0.05 False Discovery Rate value for each epitope). Because CDR3β sequences are more unique than CDR3α, more CDR3β than CDR3α paratopes were enriched across all epitopes (Supplementary Table 5). Interestingly, only one particular paratope motif, SQG, standed out as strongly enriched in narcolepsy versus controls in pHA_273-287_-tetramer positive cells (Supplementary Table 5). This motif is unique to TRBV4-2 and TRBV4-3 and reflected a significant enrichment of a subset of diverse SQG positive unique CDR3β in 6 narcolepsy patients versus 2 controls. Extended Data Figure 4 also show preferential usage of specific aminoacids in tetramers versus control sequences at various length, an analysis that was not revealing beyond the GLIPH analysis. We note that in comparison to bulk sequencing of total CD4+ memory cells, tetramer specifc CDR3β sequences had frequencies ranging from undetectable (<1/10^7^) to 1 for 10^5^ in the same corresponding subject.

### Commonly used CDR3α and CDR3β segments observed with pHA_273-287_ and NP_17-31_ tetramers

We next analyzed sharing of CDR3α, CDR3β and CDR3α/β pairs across patients and controls, with all CDR3 found at least twice across subjects (Figure 3). Not surprisingly, sharing of CDR3α was more frequent than sharing of CDR3β and exact CDR3α/β pairs were rarely seen across subjects. Sharing for viral epitope pHA_273-287_ and NP_17-31_ tetramers was as expected from usage preference reported in Extended Data Figure 3 (for example TRAJ23 for pHA_273-287_ CDR3α CAVSPGNNQGGKLIF, CIVRVLTNQGGKLIF, CIVRVATNQGGKLIF and CAAPYNQGGKLIF). Most commonly used segments in most enriched clones were TRAV8-6, TRAJ34, TRBV7-9 and TRBV4-2 for NP_17-31_. Usage diversity for pHA_273-287_ was biased toward TRAV13-1, TRAV17 and TRBV4-2, with more limited sharing of TRBV for this epitope. We also noted frequent usage of TRAV26-1 CDR3α across all epitopes; DQ0602 is a strong transQTL of TRAV26-1 segment usage, probably because it interacts through its CDR1α with DQ0602^45^.

### HCRT_NH2_ tetramers most frequently use J24 positive CDR3α CAVETDSWGKLQF

As mentioned above, TRAJ24, TRAJ28 and TRBV 4-2 are of special interest as genetic QTL locations modulating usage of these segments are associated with narcolepsy. While TRAJ28 did not show any particular over representation in our data, we observed extensive sharing of TRAJ24 and TRBV4-2. The most striking finding was extensive sharing of TRAJ24 CDR3α TRAV2-CAVETDSWGKLQF-TRAJ24 patients and controls with HCRT_54-66-NH2_ and HCRT_86-97-NH2_ respectively (Figure 3c and d). As TRAJ24 is a low usage segment (0.87%), this is highly significant. Supplementary Tables 6 and 7 report on all TRAJ24 and TRBV4-2 positive clones found in all cultures, respectively. We focused on clones containing TRAJ24 CDR3α CAVETDSWGKLQF the major CDR3α shared across patients and controls following HCRT_NH2_ stimulation (Figure 3). All these clones also use TRAV2. As can be seen, the CAVETDSWGKLQF CDR3α generally occurs in the context of diverse CDR3βs containing TRBV15, TRBV2, TRBV29-1, TRBV9 in controls and TRBV15, TRBV3-1, TRBV7-2 and TRBV20-1 in patients. Only TRAV15-CATSSGGGGKAYGYTF CDR3β was shared by 2 patients and one control following HCRT_NH2_ culture and tetramer isolation. Of note, CAVETDSWGKFQF was not observed in any subject, control or T1N. This suggests that TCRβ is less important for interaction of these TCR α/β with HCRT_54-66-NH2_ and HCRT_86-97-NH2_ when presented by DQ0602.

### Single cell sorting and TCR sequencing of INFγ positive, Pandemrix^®^ reactive CD4^+^ T cells confirms the importance of pHA_273-287_ and NP_17-31_ as immunodominant peptides of the HLA class II response to Pandemrix^®^ and suggest mimicry with HCRT CD4^+^ responses

CarboxyFluorescein Succinimidyl Ester (CFSE)-tagged PBMCs from 8 patients and 8 controls were expanded (overlapping with the sample above) in the presence of Pandemrix^®^ in a 7-day period following which CFSE-low CD4^+^ cells were sorted and expanded further for a 4day period and then stimulated with autologous monocytes prepulsed with Pandemrix^®^, following which single cell sequencing of INFγ-positive (Pandemrix^®^-reactive), CFSE low (having divided in the presence of Pandemrix^®^ during the culture) CD4^+^ cells was performed. This experimental design amplifies all Major Histocompatibility Complex (MHC) Class II mediated responses, with DQ0602 mediated responses likely in the minority, notably because DRB1 responses are typically dominant among Class II responses. To limit this problem, one patient repeated twice was also cultured with Pandemrix^®^ in the presence of anti DR and anti DP antibodies. In this case, presentation and reactivity should be limited or enriched for DQ response (DQ0602 and the other trans located allele). Upon completion of these experiments, CDR3 TCRα and TCRβ sequences were matched with tetramer-derived sequences to search for commonality (Supplementary Table 8). Strikingly, we found that pHA_273-287_ and NP_17-31_ tetramer sequences, in many cases sharing both TCRα and β chains were present in these cultures, most notably in the cultures than had been blocked by anti DR and DP antibodies. Interestingly, we also found a significant number of sequences shared with HCRT_54-66-NH2_, HCRT_86-97-NH2_ tetramer sequences suggesting mimicry with one or several unknown peptide variants in Pandemrix^®^. As a control, no sharing was found with single sorted CD8^+^ cell TCR sequences reactive to Pandemrix^®^ were isolated using a similar procedure (data not shown).

### Selected pHA_273-287_ associated CDR3β are found in HCRT_NH2_ J24 positive CDR3α CAVETDSWGKLQF positive tetramers, with a major node using TRBV4-2 CDR3β CASSQETQGRNYGYTF across HCRT_NH2_ and pHA_273-287_ tetramers. Such connections are not found with NP_17-31_ positive tetramers

In Figure 4, paired CRD3α and CRD3β found in various tetramers across antigens (from Supplementary table 4) were plotted as a network plot, with the thickness of connecting lines reflecting number of clones found with each denoted heterodimer. Clones found only once across epitopes within the same patient were removed to avoid the possibility of contamination. Tables in Figure 4b-4d indicate key used segments in this connectome. As can be seen, a clearly separated cluster that uses TRBV7-9, TRAV8-6, TRAV6, TRAJ34, and TRBV4-2 was formed with NP_17-31_ (Figure 4d). In contrast, clusters of HCRT_NH2_ and pHA_273-287_ are tightly connected (Figure 4b and 4c), reflecting not only sequence homology between these epitopes and increased sharing of specific CDR3α and CDR3β (Supplementary Table 4) but also revealing two groups of antigen-specific heterodimers (HCRT_NH2_ and pHA_273-287_-specific) connected through a few discrete likely crossreactive nodes.

**Figure 4.**
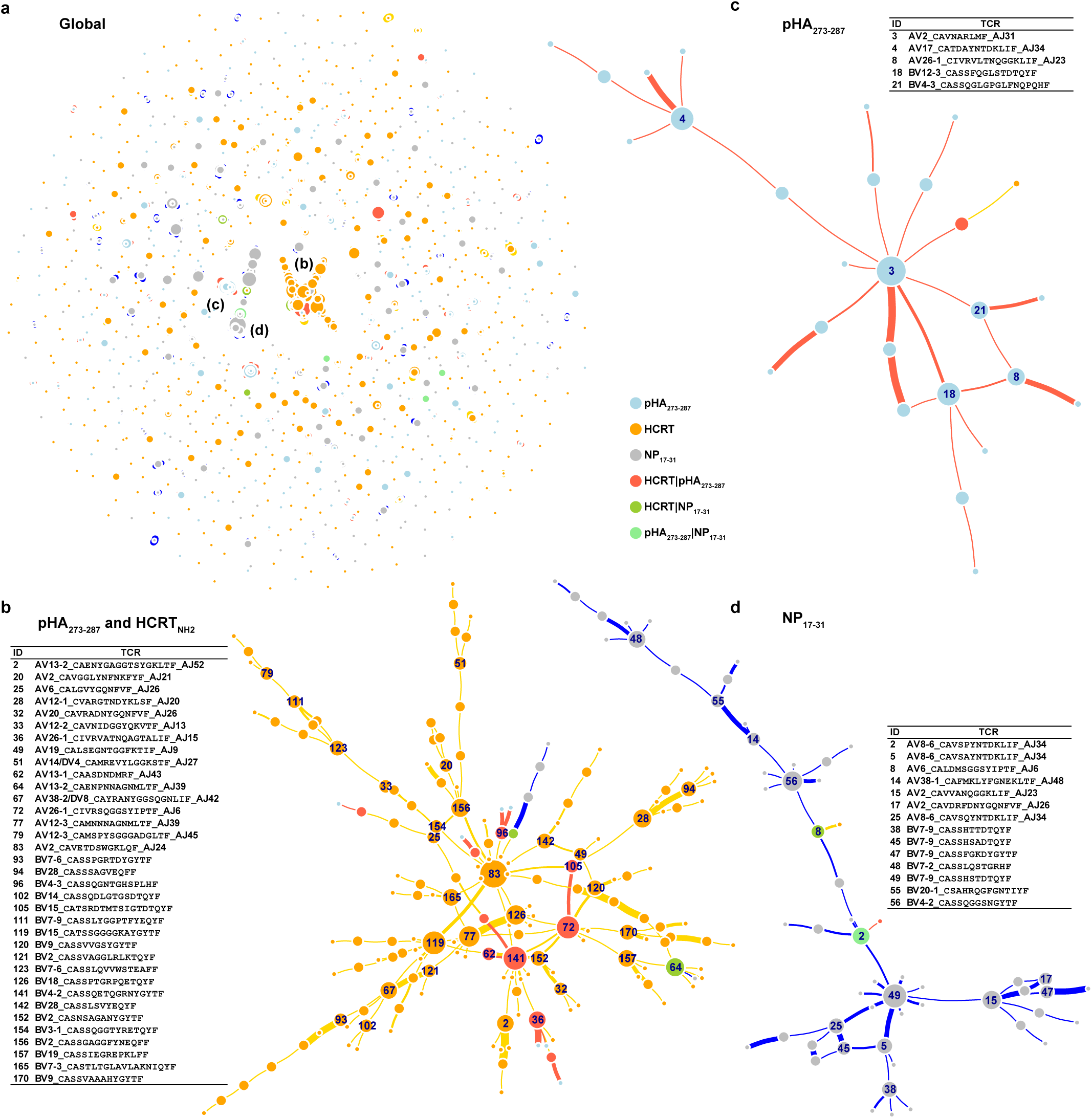
Network clustering of TCR segments. **a**. Global view of network with 3 central clusters. **b**. pHA_273-287_/HCRT_NH2_ cluster centered around TRAJ24. **c**. pHA_273-287_ only cluster. **d**. NP_17-31_ only cluster.

The HCRT_NH2_ reactive node is centered around the identified CDR3α TRAV2-CAVETDSWGKLQF-TRAJ24 (Figure 4b). As mentioned above, this TCRα chain connects with a large variety of diverse TCRα sequences, suggesting that HCRT_NH2_ may interact more primarily with TCRα. Other HCRT_NH2_ nodes, such as TRBV2 CASSGAGGFYNEQFF, TRBV3 CASSQGGTYRETQYF and TRAV12-2 CAVNIDGGYQKVTF TRAVJ23 are connected to it.

pHA_273-287_ reactive nodes are multiples and include: CDR3β TRBV4-2 CASSQETQGRNYGYTF, TRBV15 CATSSGGGGKAYGYTF, TRBV18 CASSPTGRPQETQYF, CDR3α TRAV13-2 CAENYGAGGTSYGKLTF TRAJ52 and TRAV26-1 CIVRSQGGSYIPTF TRAJ6 among others. Presumably TCR α/β containing these chains have high affinity for pHA_273-287_ in various combinations.

Most interestingly, pHA_273-287_ and HCRT_NH2_ positive nodes join in several places through less frequent heterodimers that also use the primary HCRT_NH2_ CDR3α TRAV2-CAVETDSWGKLQF-TRAJ24. These include HCRT_NH2_ reactive CDR3βs: TRBV15 CATSRDTMTSIGTDTQYF (2 clones in 2 controls), TRBV4-2 CASSQETQGRNYGYTF (one clone in one patient) and TRBV4-3 CASSQGNTGHSPLHF (one clone in one control) and TRBV4-3 CASSQGDVNYGYTF (one clone in one control)) (see supplementary Table 5 and 6). These CDR3βs are used by other heterodimers with distinct CDR3αs in the context of pHA_273-287_. Of further interest is the fact that CDR3β TRBV4-2-CASSQETQGRNYGYTF is paired with CRD3α TRAV13-1-CAASDNDMRF-TRAJ43 (25 occurrences in one subject) in the context of pHA_273-287,_ indicating it is an important heterodimer for this response. Of note, one direct connection between TRAV2 CAVETDSWGKLQF TRAJ24 and TRBV20-1 CSASPLGRGTEAF (17) primarily used by NP_17-31_ is also present, suggesting that some limited mimicry could also occur with NP_17-31_. Other connections are occurring through CDR3α, which are generally less antigen specific.

## Discussion

In this work, we screened two potential autoantigens for DQ0602 binding and T cell reactivity using tetramer staining of PBMCs of T1N cases and controls. Hypocretin and RFX4 were selected because of their high enrichment in hypocretin cell^44^. RFX4 is interesting because of its primary expression in testis and its involvement in midline brain development^48^. Furthermore, the only known established autoimmune disease with secondary hypocretin cell loss is anti-Ma2 encephalitis, a disease associated with seminomas and CD8^+^ T-mediated destruction of hypocretin cells^49^. We found only limited TCR reactivity directed towards any segment of the native HCRT peptide sequence (or RFX4), but strong reactivity to the N-Amidated, post-translationally modified (PTM) C-terminal end of hypocretin-1 and 2 (denoted HCRT_54-66-NH2_ and HCRT_86-97-NH2_ or, as a group, as HCRTNH2) (Figure 1). The two peptides, also called orexin-A and orexin –B, are homologous peptides that differ only in their C-terminal amidated residue, L-_NH2_ for orexin-A and M-_NH2_ for orexin B.. These are secreted by hypocretin neurons after undergoing amidation of their C-terminal end, a PTM necessary for biological activity^50^. The observation of higher reactivity of HCRT_NH2_ extends other observations made in other autoimmune diseases, such as Rheumatoid Arthritis, Celiac disease and Type 1 Diabetes^51^, indicating that PTMs are frequently present in autoantigens. The current explanation is that PTMs do not occur sufficiently in the thymus during central tolerance establishment^52^, and thus increase the risk of autoimmunity.

Relevant to T1N, we also found that patients have higher numbers of CD4^+^ T cells binding HCRT_54-66-NH2_ when presented by DQ0602, the major HLA allele associated with T1N (97% versus 25% in controls) (Figure 2). Recent onset cases also showed significantly higher numbers of T cells recognizing HCRT_86-97-NH2_, a peptide differing only in its C-terminal amino-acid from HCRT_54-66-NH2_. These results indicate that autoreactivity to amidated HCRT_NH2_ fragments is common in both patients and controls but occurs at higher levels in patients. These findings are consistent with other publications reporting occasional cross-reactivity to HCRT in both patients and controls, although these studies used ELISpots in a small number of samples, finding no differences by disease status^19,20^. The findings above could reflect causality, cross-reactivity secondary to hypocretin cell loss via another mechanism, or an epiphenomenon. Strikingly however, not only was sharing of CDR3 sequences found across patients (as recently reported for gluten-reactive T cells in Celiac Disease^53^) but the top CDR3α CAVETDSWGKLQF involved in the recognition of HCRT_NH2_ in controls and patients uses TRAJ24 (Figure 3), a segment marked by SNPs rs1154155G and rs1483979 strongly associated with T1N risk^6^. This genetic association is the strongest association after HLA across all ethnicity^1–6^ and as TRAJ24 is normally only used rarely (0.8% of repertoire)^29^, the result must indicate causality. In addition to that, we found common sharing of CDR3β TRBV4-2 CASSQETQGRNYGYTF in cases and controls with HCRT_NH2_ and pHA_273-287_ across 4 subjects (Figure 3, Supplementary Table 4). This is likely to be functionally significant as TRVB4-2 is also rarely used (0.7% of repertoire)^29^, and expression of this segment is linked with narcolepsy-associated SNP rs1008599^6,29,45^.

Considering these associations, our data indicate that T1N is caused by autoimmunity to HCRT_NH2_ either through dysregulation of regulatory T cells or through increased pathogenicity of the TRAJ24F allele. Because controls have the same CDR3α however, it is likely not sufficient in itself to escalate to full hypocretin cell loss and establishment of the T1N phenotype. Additional phenotyping of autoreactive cells carrying CDR3α CAVETDSWGKLQF will be required in order to answer these questions, together with CD8^+^ T cell studies. Genetic^29^ and animal studies^30^ indeed suggest a CD4^+^ helper CD8^+^ cytotoxic model as the most likely to explain hypocretin cell loss in T1N, although involvement of cytotoxic CD4^+^ T cells is also possible.

Furhermore, we demonstrate that T1N is associated with differential reactivity to at least two flu peptides, pHA_273-287_ and NP_17-31_, derived from X-179A, the reassortant strain used in Pandemrix^®^ (Figure 2). This vaccine has been linked with T1N onset in Northern Europe^34^, however the latter correlation varies across countries^34^, suggesting interactions with other factors. Of particular interest was the fact that one peptide, pHA_273-287_, is specific of 2009H1N1, while another peptide NP_17-31,_ is specific of the vaccine backbone strain PR8, an old seasonal H1N1 strain originally derived from 1918H1N1. The NP_17-31_ sequence is representative of seasonal H1N1 prior to the 2009 pandemic flu, a strain currently circulating at a lower frequency. Whether or not involved in mimicry, our data also suggest that flu epitopes derived from multiple H1N1 strains (pH1N1, PR8-like) may be needed to favour immune response towards T1N immunity, and that overall homology between HCRT and flu sequences may be of greater importance. If this was the case, a particular history of coincident flu infections, together with specific past exposure, could preferentially trigger T1N, explaining why the H1N12009 pandemic may have led to a more compelling increase of T1N in China^10,33^ as supposed to other countries. Influenza-A creates low fidelity copies and mutates in vivo during the course of infections or in specific vaccines, which could also contribute to rare mutated epitope exposure in some cases^35,54^. In this context, Pandemrix^®^ vaccination coincidental with pH1N1 pandemic or other strains may have been a perfect storm in some countries. The confluence of exposure to multiple epitopes could also explain why the administration of Arepanrix^®^, an adjuvanted vaccine similar to Pandemrix^®^ which was used in Canada, has not exhibited a strong link to T1N^55^.

pHA_273-287_ has significant homology with HCRT_54-66_ and HCRT_86-97_ sequences and their binding repertoire^18^, making it a suitable mimicry candidate with HCRT_NH2_ (Figure 3). This is reflected by increased sharing of CDR3α and CDR3β sequences in the corresponding tetramers, that was high within HCRT_NH2_ peptides, substantial between HCRT_NH2_ and pHA_273-287_ but almost absent between HCRT_NH2_ and NP_17-31_ or between pHA_273-287_ and NP_17-31_ (Supplementary Table 4). This almost linear relationship suggests the more homology the mimic has with the autoantigen sequence, the most likely similar TCR sequences are recruited by crossreactivity. Surprisingly however, the number of epitope-specific GLIPH paratopes shared between sequences was similar. We also tried to examine whether TCR paratope motifs common to HCRT_NH2_ and pHA_273-287_ recognition were more abundant in HCRT_NH2_ positive tetramers from patients versus controls, however no compelling observations were made (data not shown).

Most revealing was a network analysis examining CDR3α and CDR3β sharing across pHA_273-287_, NP_17-31_ and HCRT_NH2_ tetramers (Figure 4). In this analysis, clearly separate clusters of connected CDR3α-CDR3β heterodimers emerged for NP_17-31_ tetramers and a subset of pHA_273-287_ heterodimers. In addition to this, a large HCRT_NH2_ tetramer cluster centered around CDR3α TRAV2 CAVETDSWGKLQF TRAJ24 was formed, but it was also connected to two strongly expressed pHA_273-287_ heterodimer nodes through HCRT_NH2_ CDR3β TRBV4-2 CASSQETQGRNYGYTF in one patient, and TRBV15 CATSRDTMTSIGTDTQYF, TRBV4-3 CASSQGNTGHSPLHF or CASSQGDVNYGYTF in a few controls (see supplementary Table 5 and 6 and Figure 4). This result is remarkable as narcolepsy is genetically associated with SNPs that regulate usage of two of these three segments, TRAJ24 and TRBV4-2, suggesting causality. For the other two connections, TRBV15 and TRBV4-3, examination of Quantitative Ttrait Loci (QTL) effects for these segments did not reveal any genomic effect^29^ unlike the well-established TRBV4-2 effect^29,45^, thus functional connection through genetic association could not be tested. Of particular interest is the size of the TRAV26-1 CIVRSQGGSYIPTF CDR3α node, likely explained by the fact that TRAV26-1 is a transQTL for DQ0602, reflecting direct CDR1/2 interactions of TRAV26-1 with the DQ0602 molecule, notably DQα, the most specific portion of the DQ molecule^45^

In the case of TRBV4-2, the connection is direct, as narcolepsy-associated SNP rs1008599A increases usage of this segment, and thus presumably the risk of the crossreactive CDR3α TRAV2 CAVETDSWGKLQF TRAJ24 / CDR3β TRBV4-2 CASSQETQGRNYGYTF heterodimer to occur. For TRAJ24, the connection is more complex as the genetic signal encompasses several SNPs (notably rs1154155G and rs1483979C), one of which changes an aminoacid within TRAJ24, so that its terminal CDR3α end region is GKFQF instead of GKLQF (L is the most common allele in Caucasians). In addition to this effect, narcolepsy-associated SNPs also decrease TRAJ24 expression and increase TRAJ28 expression (a segment not involved in our dataset). In our dataset, the major narcolepsy-associated CDR3α segment observed is TRAV2 CAVETDSWGKLQF TRAJ24, as segment containing the L allele that is protective for narcolepsy. TRAV2 CAVETDSWGKFQF TRAJ24 was not observed. This could be explained by the fact TRAJ24F is expressed at about half the level of TRAJ24L^29^ and is only present in 30% of our patients tested, most of whom are J24 L/L homozygous (the most frequent combination in Caucasians). Alternatively complex effects involving T regulatory cells with these TCR alleles may be involved. Testing of more patients carrying rs1483979C may reveal CAVETDSWGKFQF or other TRAJ24F containing segment(s) with higher HCRT_NH2_ and pHA_273-287_ crossreactivity. Supplementary Table 6 shows examples of possible TRAJ24F associated HCRT_NH2_ sequences, such as TRAV2 CAFTTDSWGKFQF TRAJ24 (10 occurences in a recent onset case) but these occurences were too few to allow any conclusion to be drawn.

An interesting aspect of this data is the strong association of HCRT_NH2_ reactivity to a specific CDR3α segment, TRAV2 CAVETDSWGKLQF TRAJ24, in association with a diversity of CDR3βs. Interestingly, if normally docked, the most conserved N-terminal end portion of HCRT_NH2 NH2_-GNHAAGILTL/M_-CO-NH2_ is expected to interact with CDR3α, whereas the more variable amidated C-terminal end should contact CDR3 β possibly contributing to this phenomena. It may be that HCRT amidation induces a broad cross reactivity via CDR3α contacts without as much of a need for tight CDR3βcontacts. In this scenario, flu responses, generally more specifically driven by CDR3β contacts^47,56^ as most usual epitopes, could occasionally amplify a rare TCR heterodimer containing CDR3β TRBV15 CATSRDTMTSIGTDTQYF, TRBV4-2 CASSQETQGRNYGYTF, TRBV4-3 CASSQGNTGHSPLHF or TRBV4-3 CASSQGDVNYGYTF coupled with TRAV2 CAVETDSWGKLQF TRAJ24 (nodes connecting pHA_273-287_ and HCRT_NH2_ networks in Figure 4b), inducing autoimmunity via cross-reactivity to HCRT_NH2_. Recent analysis of ∼80 available TCR-pMHC structures (26 MHC class II) has shown that whereas there is always at least one CDR3β contact with pMHC, there are cases where no CDR3α contact occurs^47,57^. Reverse peptide docking has also been reported in some cases^57^ with CDR3β and not CDR3α contacting peptide residues. Similarly, an autoreactive TCR–MHC class II complex found extreme amino-terminal positioning of the TCR over the antigen-binding platform of the MHC molecule, suggesting a link between atypical TCR docking modes and autoreactivity†^58^, althrough this has not been found in other autoreactive structures. We hypothesise that once a mostly CDR3β driven flu response has TCRα-“hitchhiked” a rare autoreactive clone containing the culprit TRAJ24 CDR3α sequence, HCRT_NH2_ released by damaged hypocretin cells would then amplify a more diverse array of TRAJ24 CDR3α/β response involving other CDR3β with better contacts with HCRT_NH2_ such as TRBV7-2 CASSYDRGSKPQHF, TRBV3-1 CASSQGGTYRETQYF, TRBV20-1 CSASPLGRGTEAFF, TRBV15 CATSSGGGGKAYGYTF, TRBV2 CASSGAGGFYNEQF, TRBV29-1 CSVVSTSASYNEQFF or TRBV9 CASSGTLQGTGGYTF (Supplementary Table 6). Additional studies involving transfection of these heterodimers in TCR reporter-containing TCR deficient Jurkat cell lines, with activation with cognate pHA_273-287_ and HCRT_NH2_ ligands displayed by DQ0602 will be needed to confirm these hypotheses. Cross cultures and structural studies of individual clones would then naturally follow.

A number of molecular mechanisms have been suggested to explain HLA association in autoimmunity^51^. In narcolepsy, major factors likely involve tolerance escape due to PTMs of the hypocretin autoantigen and crossreactivity with specific flu sequences which may be favored by a primary role of CDR3α in binding this particular autoantigen (a variation of the hotspot binding molecular mimicry hypothesis^51^). Although this study is the first to link specific TCR heterodimers to an autoimmune disease through TCR tetramer sequencing and mendelian randomization, many questions remain un-answered. These include: What is the role of the narcolepsy-associated TRAJ24F allele in disease predisposition? Which flu epitopes besides pHA_273-287_ are involved, notably prior to 2009? What is the role of NP_17-31_, as minimal or no evidence for mimicry was found with this peptide? What is the role of regulatory T cells and of occasional autoreactivity to non-amidated HCRT or RFX4 fragments? What is the cellular phenotype of the identified autoreactive cells? What portion of normal subjects have crossreactive CDR3α/β-containing T cells in their naïve compartments? Is the presence of TRAV2 CAVETDSWGKLQF TRAJ24 with TRBV4-2 CASSQETQGRNYGYTF in a patient a rare, stochastic occurrence? More extensive tetramer sequencing studies using bar coded tetramers will be needed to fully understand the spectrum of cross reactivity of flu and HCRT_NH2_ autoreactive CD4^+^ T cells involved in T1N. Most importantly, HCRT_NH2_ autoreactivity appears to be necessary, but not sufficient to induce a full hypocretin neuronal loss and T1N since controls also exhibit some autoreactivity and this remains to be explained. Involvement of regulatory T cells, of other autoantigen(s) and of CD8^+^ T cells could complete a complex pathophysiological picture, which may serve as a guide to understanding other autoimmune diseases with more complex genetic associations.

## Methods

### Ethics Statement

This study was reviewed and approved by the Stanford University Institutional Review Board (Protocol # 14325, Registration # 5136). Informed consent was obtained from each participant.

### Participants

All patients with narcolepsy (n = 35; 45.7% male; mean age ± standard error of the mean (SEM), 19.0 ± 1.6 years; range, 6 to 53 years) had cataplexy with an onset age of 18.9 ± 1.7 (mean ± SEM) years (ranging from 6 to 53 years) and meet criteria for International Classification of Sleep Disorders 3 (ICSD 3) for type 1 narcolepsy. Cerebrospinal fluid (CSF) hypocretin-1 levels were tested from available 7 patients and all had low level (mean ± SEM, 28.04 ± 13.51 pg/ml, see Supplementary Table 9). 16 out of 35 (45.7%) patients had onset post Pandemrix vaccination, less than one year following vaccination (all in 2009-2010, except for one case vaccinated with Pandemrix in the following season). Controls (n = 22; 59.1% male; mean age ± SEM, 23.3 ± 2.9 years; range, 10 to 57 years) were unrelated subjects, healthy twin or vaccinated siblings. 11 out of 22 (50.0%) controls were vaccinated with Pandemrix. All patients and controls were DQB1*06:02 positive. Peripheral blood mononuclear cells (PBMCs) were collected by apheresis at the Stanford blood center or through phlebotomy, followed by ficoll isolation and storage in liquid nitrogen until use.

### Vaccine

The Pandemrix vaccine (A/California/7/2009 (H1N1) NYMC X-179A monovalent bulk (inactivated, sterile)) (Batch# AFLSFDA280) used throughout the study was manufactured with HA content at 139 μg/ml (determined with single radial diffusion (SRD)) by GlaxoSmithKline (GSK) Dresden in January 2010. This batch has been used during the 2009-2010 vaccination campaign in Europe. Summary of Pandemrix characteristics can be found in European medicines agency document (http://www.ema.europa.eu/docs/en_GB/document_library/Other/2010/05/WC500091295.pdf).

### Peptides

Overlapping 15-mer peptides (11-amino acid overlap) covering full length of hemagglutinin (HA), neuraminidase (NA), polymerase (basic) protein 1 (PB1), prepro-hyprocretin (HCRT, including post translational modification), 23 variants of nucleoprotein (NP) derived from A/California/07/2009 and A/Puerto Rico/8/1934, and 6 isoforms of regulatory factor X4 (RFX4) (Supplementary Table 1) were synthesized with >90% purity at GenScript NJ and dissolved in dimethyl sulfoxide (DMSO) at a stock concentration of 5 mM or 10 mM as applicable.

### Construction of DQ0602

Generation of soluble class II major histocompatibility complex (MHC) was previously described^59^. For DQ0602, truncated coding sequences (CDS) (deletion of signal peptide and transmembrane domain) of human leukocyte antigen (HLA)-DQB1*06:02 (residues 33-230 of NP_002114.3) and HLA-DQA1*01:02 (residues 24-217 of NP_002113.2) were amplified with specific primers (DQB1-F: 5’-TATGGATCCCGGGGCCGGCCCAGTGTCCAAGAT-3’, DQB1-R: 5’-TATATGAATTCGCCACCTCTAGACTTGGACTGAGCGCTCTCGC-3’; DQA1-F: 5’-TATGGATCCCGAGGATATCGTCGCCGACCATGTCGC-3’, DQA1-R: 5’-TATATGAATTCGCCACCTCTAGACTCTGTGAGTTCGCTCATAG-3’) using plasmid DQB1-0602 CLIPmaster (gift from the Emory University NIH tetramer core facility) as a template, respectively. The DQB1 together with class II-associated invariant chain peptide (CLIP) and thrombin cleavage site and DQA1 were then next cloned downstream of a GP67A signal sequence and upstream of a 6*His tag in baculovirus expression vectors 1G4TCRβ and 1G4TCRα^60^ (gifts from Garcia lab at Stanford) after double digestion (BamHI/EcoRI), respectively, resulting in 2 chimeric cassettes: CLIP-thrombin cleavage site-DQB1-acidic leucine zipper-biotinylation site-6*His tag and DQA1-basic leucine zipper-6*His tag. As above, HLA-DMA (residues 1-231 of NM_006120.3) and HLA-DMB (residues 1-218 of NM_002118.4) were cloned into plasmids 1G4TCRα and 1G4TCRβ^60^ after double digestion (XmaI/EcoRI), respectively. Human cDNA library was used as a template for HLA-DM (DM) with specific primers (DMA-F1: 5’-TCCCCCCGGGATGGGTCATGAACAGAACCAAG-3’, DMA-R1: 5’-CCGGAATTCCACATTCTCCAGCAGATCTGAGG-3’; DMB-F1: 5’-TCCCCCCGGGATGATCACATTCCTGCCGC-3’, DMB-R1: 5’-CCGGAATTCCTTCAGGGTCTGCATGGGG-3’).

### Peptide exchange

Purification of monomer class II MHC was previously described^60,61^. Transfection and biotinylation of DQ0602 were conducted using BestBac^®^ 2.0 (Cat# 91-002, Expression Systems) and BirA (Cat# BirA500, Avidity) according to manufacturers’ instructions, respectively. Biotinylated DQ0602 (Bio-DQ0602) was isolated by fast protein liquid chromatography (FPLC) at Biomaterials and Advanced Drug Delivery (BioADD) laboratory at Stanford. Peptide exchange^62^ was validated with 3 2,4-dinitrophenyl (DNP)-conjugated peptides: Epstein-Barr virus (EBV) nuclear antigen-1 (EBV_486-500_, RALLARSHVERTTDE)^63–65^, cytomegalovirus (CMV) phosphoprotein pp65 (CMV_41-54_, LLQTGIHVRVSQPS)^66^, and herpes simplex virus type 2 (HSV-2) VP16 (HSV_33-50_, PPLYATGRLSQAQLMPSP)^59,67^. Briefly, 5 μg of cleaved Bio-DQ0602 (1 mg/ml) was incubated with 0.25% octyl-β-D-glucoside, 100 mM citrate/NaH_2_PO_4_ (pH=5.2), 0.4 mg/ml DNP-peptide, and DM (molar ratio of DM:DQ0602=4:1) for 72 h at 37°C, followed by neutralization by adding 1/5 volume of 1 M Tris-HCl (pH=8.0). To analyze peptide exchange efficiency, streptavidin microspheres were added and run on BD LSRII at Stanford Shared FACS Facility (SSFF) after staining with allophycocyanin (APC) anti-HLA-DQ (Cat# 17-9881-42, eBioscience) and anti-DNP (Cat# ab24319, Abcam; isotype, mouse IgE) antibodies, followed by secondary fluorescein isothiocyanate (FITC) antimouse IgE (Cat# ab99572, Abcam) antibody.

### Generation of peptide-DQ0602 (pDQ0602) tetramer

Generation of class II tetramer was previously described^59,62^. Briefly, the neutralized reaction (above) was incubated with phycoerythrin (PE)-streptavidin (Cat# 405204, BioLegend) at 6:1 molar ratio of Bio-DQ0602:PE-streptavidin for 1 h at room temperature (RT) in dark. Extra Bio-DQ0602 was removed by streptavidin agarose using a centrifugal filter unit (REF# UFC30GV00, EMD Millipore). pDQ0602 tetramer was concentrated in phosphate buffered saline (PBS, pH=7.4) by centrifugation (REF# UFC510096, EMD Millipore).

### Pandemrix cell culture and pDQ0602 tetramer sorting

Tetramer staining was previously described^59,62,68,69^. Briefly, PBMCs were stimulated with100 ng/ml (final concentration of HA) Pandemrix in a 50 ml-culture flask (2.5×10^6^ cells/ml) or 6.25 μM peptide in a 96-well plate (1-2.5×10^6^ cells/ml) for 10 or 14 days at 37°C, 5% CO_2_. 20 IU/ml IL-2 (Cat# 14-8029-81, eBiosciences) was supplemented from day 7 in complete RPMI medium (RPMI (Cat# 61870-036, Gibco) supplemented with 10% fetal bovine serum (FBS) and 1% penicillin/streptomycin). Medium was changed every 2-4 days. Immediately before staining, pDQ0602 tetramer was spun down to remove aggregates (if any). Fresh or supernatant pDQ0602 tetramer (>10 μg/ml) was incubated with cultured PBMCs in 100 μl of complete RPMI medium for 90 min at 37°C, 5% CO_2_. Cells were then stained with a combination of Brilliant Violet 421^®^ (BV421) anti-CD3, Alexa Fluor^®^ 488 (AF488) anti-CD4, Alexa Fluor^®^ 700 (AF700) anti-CD8 and APC/Cy7 anti-CD45RO antibodies (all from BioLegend), followed by analysis on BD LSRII or sorting on BD ARIA II at SSFF. Propidium iodide (PI) was used to gate live cells. Data was analyzed using FlowJo (v10.0.8r1). For most experiments, multiple repeats were done, and if this was the case the mean of repeat observations for each subject was used in statistical analysis.

To complement tetramer data, CD4^+^ T cells that had been activated by Pandemrix were enriched utilizing an interferon-γ (IFNγ) secretion assay. Briefly, 10-20 million PBMCs were tagged with 0.5 μM carboxyfluorescein diacetate succinimidyl ester (CFSE) (REF# C34554, Life Technologies) in the presence of 100 ng/ml (HA) of Pandemrix and expanded for a 7-day period, whilst supplementing the culture as needed. On day 7, cultured PBMCs were harvested and an aliquot was phenotyped for quantification of T helper subsets as described^70^ on BD LSRII. To sort CFSE-low cells, the remaining portion was then stained with a combination of AF647 anti-CD3 (BioLegend), PE/Cy7 anti-CD4 (BioLegend), AF700 anti-CD8 (BD Biosciences) antibodies and the viability dye 7AAD (BioLegend). The CFSE-low CD4^+^ T cells, which contain the bulk of proliferating cells in response to Pandemrix stimulation, were sorted (minimum of 200,000 cells using a BD ARIA II) into sterile tubes containing advanced RPMI (Cat# 12633-020, Gibco) supplemented with 10 % FBS and 25 mM (4-2-hydroxyethyl)-1-piperazineethanesulfonic acid (HEPES). As a control, an equal number of CFSE-high cells from the CD4 gate, were sorted as described above. The sorted CD4^+^ T cells were further cultured in high dose IL-2 (500 IU/mL) for 4 days, following which autologous monocytes (CD14 pre-pulsed with 1 μg/mL Pandemrix for 4 h) were added to stimulate sorted CFSE-low versus CFSE-high CD4^+^ cells in a 1:2 ratio for a 16 h period. Following this, viable IFNγ producing CD4^+^ T cells were enriched using an IFNγ secretion assay (Miltenyi Biotec) with the two different cell types (CFSE-low & CFSE-high). These were then single cell sorted for T cell receptor (TCR) sequencing (TCRseq) using a BD ARIA II^71^. In subset of individuals (n=2) the pulsing of monocytes with Pandemrix was performed in the presence of 10 μg/mL each of anti-HLA-DP (REF# H260, Leinco Technologies) and anti-HLA-DQ (Cat# ab23632, Abcam) antibodies.

### Competition binding assay

Peptide binding assay was conducted as previously conducted^59,72^. Briefly, 25 nM cleaved DQ0602 was incubated with 100 nM DM, 1 µM biotin-conjugated EBV epitope (Bio-EBV_486-500_, Bio-(GGG)RALLARSHVERTTDE), and 40 µM competitor peptide in reaction buffer (100 mM acetate, pH=4.6, 150 mM NaCl, 1% BSA, 0.5% Nonidet P-40, 0.1% NaN_3_) with duplicate for 72 h at 37°C. The reaction was quenched by adding two volumes of neutralization buffer (100 mM Tris-HCl, pH=8.6, 150 mM NaCl, 1% BSA, 0.5 Nonidet P-40, 0.1% NaN_3_), followed by incubation with monoclonal anti-DQ (SPV-L3) antibody (Cat# BNUM0200-50, Biotium) (1:400 in 100 mM carbonate-bicarbonate buffer, pH=9.5) in a high binding 96-well plate (REF# 9018, Corning). DELFIA^®^ time-resolved fluorescence (TRF) intensity was detected using a Tecan Infinite^®^ M1000 after successive incubation with 100 µl/well Europium (Eu)-labelled streptavidin (Cat# 1244-360, PerkinElmer) (1:1000, 1% BSA in PBS) and 100 µl/well enhancement solution (Cat# 1244-105, PerkinElmer). Plate was washed 5 times with 300 µl/well wash buffer (0.05% Tween-20 in PBS) to remove nonspecific binding.

### Single cell TCR sequencing

Single cell paired TCR sequencing was performed as described_71,73_. Briefly, single cells of CD3^+^CD4^+^CD8^-^Tetramer^+^ or IFNγ^+^CD4^+^ were sorted directly into OneStep^®^ reverse transcription polymerase chain reaction (RT-PCR) buffer (Cat# 210215, Qiagen) in a 96-well plate, following which the plates were frozen at −80°C until assayed. Paired TCR CDR3 sequences were obtained by a series of 3 nested PCR reactions. For the first reaction, reverse transcription and pre-amplification were performed with multiplex PCR in a 20 μl of reaction using multiple Vα/Vβ and Cα/Cβ region primers and using the following cycling condition: 50°C 30 min; 95°C 15 min; 94°C 30 s, 62°C 1 min, 72°C 1 min × 25 cycles; 72 °C 5 min; 4 °C 10 min. The second PCR was performed with HotStarTaq^®^ DNA polymerase (Cat# 203205, Qiagen) in a 20 μl of reaction using 1 μl of the first reaction as template and the following cycling condition: 95°C 15 min; 94°C 30 s, 64°C 1 min, 72°C 1 min × 25 cycles (TCR); 72 °C 5 min; 4 °C 10 min. The third PCR incorporated barcodes to enable sequencing on the Illumina^®^ MiSeq platform and was performed in a 20 μl of reaction using 1 μl of the second PCR product as template. Combined PCR products were run on a 1.2% agarose gel. DNA of around 350 to 380 bp was purified and then sequenced on MiSeq platform for a pair-ended 2×250 run. Fastq files were analyzed with a customized TCRseq pipeline at the Human Immune Monitoring Center (HIMC) at Stanford.

### TCR Bulk sequencing

TCRα and β repertoires of CD4^+^ memory T cells isolated from PBMCs were amplified from total ribonucleic acid (RNA) by ARM-PCR procedure as previously described^74^ and then sequenced by 2×101 HiSeq 2000 and 2×151 HiSeq 4000 at the Stanford Genome Sequencing Service Center. Data analysis was briefly as follows. The quality of raw fastq files was assessed using FASTQC (http://www.bioinformatics.babraham.ac.uk/projects/fastqc). FLASH was used to merge the paired end reads^75^. BLAST databases were constructed from IMGT TCRα/β sequences using local blast binaries^76^. The fastq file was converted into fasta and simultaneously processed into equal chunks of fasta records which were then processed using custom python-based analysis pipeline implemented on the Stanford University Genomics Cluster. Each fasta chunk was used to query the BLAST database and identify the TCR C, J & V regions. The CDR3s were then extracted and their frequency along with usage of V, J families computed.

### Statistical analyses

Statistical comparisons were performed using two-tailed Mann-Whitney U tests, paired Wilcoxon non parametric tests or Fisher’s exact tests whenever most appropriate. Data were plotted using GraphPad PRISM 5. P-value <0.05 were considered statistically significant. All data wrangling and handling was implemented with custom python scripts. R function mantelhaen.test was used to perform Mantel-Haenszel chi-squared test to assess enrichment of TCR families comparing bulk sequencing to single cell sequencing.

For network construction of Figure 4, paired CDR3α/β calls and available alternate CDR3αs for each peptide-specific T cell were pooled in a database. In this database, available CDR3α or CDR3β must pair with one or more corresponding CDR3s across the dataset or are eliminated. Each CDR3α or CDR3β was then used as a key and corresponding CDR3’s that pair with the key used as values and matched across the whole dataset using a custom python based pipeline. These matched CDR3αs and CDR3βs was used as template to construct a network object to build a global network of CDR3α and CDR3β calls across peptides and samples. A python based recursive traversal algorithm was implemented to find local networks i.e. given a CDR3α or CDR3β string, find all the connected CDR3 clusters directly or indirectly to the input CDR3α. This approach was used to extract the TRAJ24 HCRT_NH2_/ pHA_273-287_, NP_17-31_ and pHA_273-287_ local clusters of CDR3s. Node size was calculated based on how many connections a particular CDR3 accepts or sends out, while the edge or connected line thickness was based on total counts of that particular CDR3α/β pair.

### Legends to Figures

Figure 1: Results of primary tetramer screening in 10 subjects: a. Tetramer positivity in post-Pandemrix^®^ patients (dark grey, from 5 total), post-Pandemrix^®^ controls (light grey, from 4 total) and one early onset patient (onset=0.3 year prior, blue, single subject). Tetramers were built one by one using peptide exchange using a DQ0602-CLIP (see Methods). Viral protein tetramers: HA=Hemagglutinin, NA= Neuraminidase, NP= Nucleoproteins, PB1= Polymerase Basic 1. Binder sequences from influenza A/reassortant/NYMC X-179A (reference ADE2909) were primarily used. X179A is a reassortant of A/California/07/2009 (pandemic pH1N1 2009) with A/Puerto Rico/8/1934. A/Puerto Rico/8/1934, is an old pre-2009 H1N1 strain, commonly called PR8, representative of seasonal H1N1 strains prior to the 2009-2010 pandemic and still circulating at low levels. In X-179A the primary sequence of HA, NA and PB1 is from pH1N1 while the NP sequence is from PR8. Any polymorphic peptide sequence differing in NP (A/California/07/2009) between PR8 (/Puerto Rico/8/and pH1N1) that still bound to DQ0602 was also tested and is labelled with a star (others are of PR8 origin). s=epitope of identical sequence in PR8 and pH1N1. Subjects were tested with tetramers after culture with Pandemrix 100 ng/ml for 10 days. Autoantigen tetramers: HCRT= prepro-hypocretin, RFX4=all known isoforms of Regulatory Factor X4, last six peptides=post-translationally modified sequences of prepro-HCRT modified as per secreted hypocretin-1 and 2, when binding DQ0602, including N-Pyroglutamate HCRT_34-48_ plus HCRT_38-52_, HCRT_33-47_ and HCRT_37-51_ with two double disulfide bonds (DB), C-terminal amidated HCRT_54-66_ and HCRT_86-97_ denoted HCRT_54-66-NH2_ and HCRT_86-97-NH2_, as in secreted, mature hypocretin-1 and 2. Subjects were tested after a 10-day culture with 6.25 μM of each peptide. The main epitopes selected for further analysis are labelled above each bar (based on immunodominance and likely difference between narcolepsy and controls). Peptides pHA_209-223_, pHA_273-287,_ pHA_321-335_, pHA_437-451_, pNA_45-59_, pNA_217-231_, pNA_217-231_, NP_17-31_, NP_261-275s_ (shared epitope between pH1N1 and PR8), and NP_437-451-pH1N1_* are responding in 60% of subjects and are considered immunodominants in the DQ0602 response to H1N1. NP_17-31-pH1N1_ the sequence equivalent to NP_17-31_ but derived from pH1N1 instead of backbone PR8 sequence. HA69 (pHA_273-287_) does not have a strong binder in the same area for PR8. b. Distribution of positive subjects per antigen. Note that response to HA, NA and NP, the main antigens contained in Pandemrix^®^ and known to be responsive to flu infections have the large number of positive subjects (≥ 2) whereas binders from autoantigens RFX4 and HCRT only show sporadic response in a maximum of 2 subjects. c. Mean percent of subjects positive for DQ0602 binders in each proteins in post Pandemrix^®^ subjects. Note that viral peptides derived from HA, NA and NP peptides have a large number of responders (36±3%, n=55) in comparison to lower abundance protein PB1 (26±5%, n=7). The response of non-modified autoantigen sequence is lowest at 5±1%. (n=67). In contrast, the response to post translationally modified (PTM)-modified hypocretin fragment is high (40± 16%, n=6), approaching that of foreign, viral antigens (see t-test statistics in figure). d. Example of FACS staining obtained for various selected tetramers. The entire dataset of FACS staining plots is provided as Extended Data Figure 1. Note tail-like staining in many (but not all) cases with HCRT_86-97-NH2_ but not with other tetramers.

Figure 2: Results of secondary tetramer screening using pHA_273-287,_ NP_17-31,_ HCRT_54-66,_ HCRT_54-66-NH2_, HCRT_86-97,_ and HCRT_86-97-NH2_ in 35 patients and 22 controls. Tetramers were built one by one using peptide exchange using a DQ0602-CLIP (see Methods). a. Percent of subjects with pHA_273-287_ and NP_17-31_ tetramer positivity in controls versus patients. PBMCs of subjects were grown in Pandemrix^®^ or the cognate peptide for 10 days in two independent experiments, and as results were repeatable and did not differ statistically, the mean is presented in each dot. T, twins. Subjects in red are those with single cell sorting and sequencing data available. Number of subjects tested is under parenthesis. b. Percent of subjects with HCRT_54-66_/HCRT_54-66-NH2_ and HCRT_86-97_ /HCRT_86-97-NH2_ tetramer positivity in controls versus patients. PBMCs of subjects were grown with the cognate peptide for 10 or 14 days in two independent experiments, and as results were repeatable and did not differ statistically, the mean is presented in each dot. T, twins. Subjects in red are those with single cell sorting and sequencing data available. c: Example of pHA_273-287_ and NP_17-31_ tetramer staining in a few subjects (for all results, see Extended Data Figure 1). d. Example of HCRT_54-66,_ HCRT_54-66-NH2_, HCRT_86-97,_ and HCRT_86-97-NH2_ tetramer staining in a few subjects (for all results, see Extended Data Figure 1). Note the “tail-like staining” aspect of HCRT_86-97-NH2_ often found with this peptide, although occasional distinct monoclonal population can be observed

Figure 3: Shared CDR3α, CDR3β and CDR3α/β sequences for pHA_273-287_ (a), NP_17-31,_ (b), HCRT_54-66-NH2_ (c), and HCRT_86-97-NH2_ (d), across subjects. Red: number of patients with corresponding CDR3, Grey, number of controls with same CDR3 (per horizontal axis). Box numbers indicate total of times when the CDR3 occurred across patients and controls. e: pHA_273-287_, NP_17-31_, HCRT_54-66-NH2_, and HCRT_86-97-NH2_ sequence homology assessed using http://tools.iedb.org/cluster/reference/^46^ with three EBV sequences used as anchor. Note high similarity of HCRT_54-66-NH2_, and HCRT_86-97-NH2_, also share homology with pHA_273-287_^18,24^ as previously reported, but not NP_17-31._ Note extensive sharing of TRAJ24 (labelled in red) CDR3 CAVETDSWGKLQF in patients and controls with HCRT_54-66-NH2_ and HCRT_86-97-NH2_. In most cases, these CRD3α CAVETDSWGKLQF do not share the same CDR3β (see Supplementary Table 4). We also found sharing of TRBV4-2 containing CRD3β CASSQERGSYNEQFF (labelled in red) in patients with NP_17-31_ only and sharing of TRBV4-2 containing CRD3β CASSQETQGRNYGYTF (labelled in red) in pHA_273-287_ and HCRT_NH2_ associated tetramers although these do not share a similar CRD3α (see supplementary Table 4).

Figure 4:. Network clustering of TCR segments obtained through tetramer sequencing: (a) global view of the generated network with its 3 central clusters; (b) pHA_273-287_/HCRT_NH2_ cluster centered around TRAJ24; (c) pHA_273-287_ specific cluster; and (d) NP_17-31_ specific cluster. Tree formation was computed using a partional recursive clustering elimination algorithm implemented using Python. Each node represents a CDR3α or a CDR3β segment, with size proportional to occurrences. Lines connecting these clusters represent heterodimers, with thickness proportional to clone number. Color coding corresponds to type of tetramer (pHA_273-287_, HCRT_NH2_ or NP_17-31_) yielding specific clones. Note that Cluster (b), primarily derived from HCRT_NH2_ tetramers and centered around CDR3α TRAV2_CAVETDSWGKLQF_TRAJ24, is hybrid, containing not only CDR3β recruited by HCRT_NH2_ but also CDR3β chains more frequently recruited by pHA_273-287_ and occasionally recruited by HCRT_NH2_ TRBV4-2_CASSQETQGRNYGYTF (141), TRBV15_CATSRDTMTSIGTDTQYF (105), TRBV4-3_CASSQGNTGHSPLHF (96) and TRBV4-3_CASSQGDVNYGYTF (10) (see supplementary Table 5 and 6). One small direct connection between TRAV2_CAVETDSWGKLQF_TRAJ24 and TRBV20-1_CSASPLGRGTEAF (17) primarily used by NP_17-31_ is also present, suggesting that some limited mimicry may also occur with NP_17-31_. Abundantly expressed TRAV26-1_CIVRSQGGSYIPTF_TRAJ6 is a CDR3α and is indirectly linked. Note also NP_17-31_-CDR3α TRAV13-2_CAENPNNAGNMLTF_AJ39 (64), a possible link to NP_17-31_ but through a CDR3α as CDR3α are more likely to be shared by chance and this is peripheral to the network, significance is uncertain.

## Acknowledgments

The study was funded by gifts from Wake Up Narcolepsy, Jazz Pharmaceutical and individual patients to Stanford University. Some samples have been gathered by defunct NIH grant P50-NS-23724 to Emmanuel Mignot (ended in 2016). Bulk Sequencing and some sample collection has been funded by a prior grant from Glaxo Smith Kline (GSK), supervised by the European Medical Agency (ended in 2014), with corresponding data reported in Ollila et al.^29^. Note that none of the current authors have been involved in the generation of fraudulent ELISpot results reported in the retracted De la Herran-Arita et al. publication^18,24^, as determined by a review of Stanford’s internal investigation report by NIH Office of Research Integrity. Dr. Mignot occasionally consults and has received contracts from Jazz Pharmaceuticals and is and has been a Principal Investigator on clinical trials using sodium oxybate and Solriamfetol, Jazz Pharmaceutical products, in the treatment of narcolepsy; none of these have any scientific relationship to the study. This work used the Genome Sequencing Service Center by Stanford Center for Genomics and Personalized Medicine Sequencing Center, supported by the grant award NIH S10OD020141. We thank Dr. K. Christopher Garcia for sharing plasmids and insect cell lines and NIH tetramer core facility for sharing DQA1*01:02/DQB1*0602 expression construct. We thank Wenchao Sun from Stanford Biomaterials and Advanced Drug Delivery Lab (BioADD) for HPLC running and Stanford FACS facility for providing analyzers and sorters and technical support.

## Supporting information

Supplementary Table 1. Screening of HA, NA, NP, PB1, HCRT and RFX4 overlapping peptides for DQ0602 binding.

Supplementary Table 2. DQ0602 registers of main peptides and sequences used for confirmatory tetramer studies.

Supplementary Table 3. Single cell sequencing TCR segment usage preference in cases and controls.

Supplementary Table 4. Sharing of CDR3α, CDR3β and CDR3αβ pairs across epitopes/individuals.

Supplementary Table 5. Significant CDR3 motifs across peptides and individual.

Supplementary Table 6. TRAJ24 positive clones found in patients and controls with various antigens.

Supplementary Table 7. TRBV4-2 positive clones found in patients and controls with various antigens.

Supplementary Table 8. TCR sharing of tetramer with IFNγ positive cells cultured with Pandemrix.

Supplementary Table 9. Clinical characteristics and use of subjects in this study.

---

**Extended Data Figure 2.**
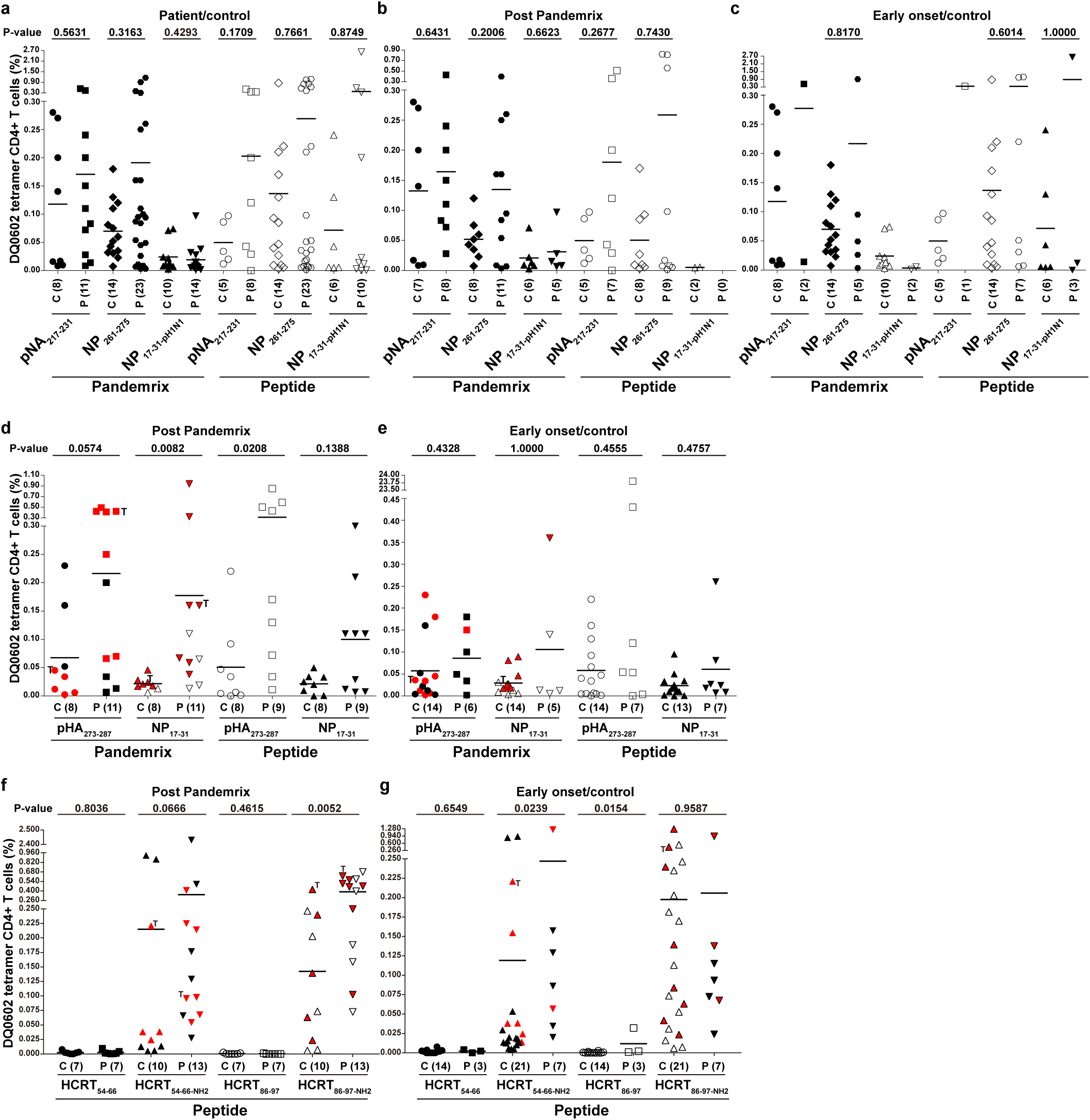
Tetramer positivity for pHA_273-287_, pNA_217-231_, NP_17-31_, NP_17-31-pH1N1_, NP_261-275_, HCRT_54-66-NH2_ and HCRT_86-97-NH2_ in patients versus controls (a), including subgroup analysis of post Pandemrix cases/controls only (b,d and f) and early onset cases versus matched controls only (c, e and g). Early onset are defined as having had onset less than one year prior to blood sample. P-value is shown on top of each panel. P, patient; C, control; T, twins; Bar indicates the mean; The number of subjects tested is shown between brackets; Red, single cell TCR sequencing. For peptide and subject infomation, see Supplementary Table 1 and 9, respectively.

**Extended Data Figure 3.**
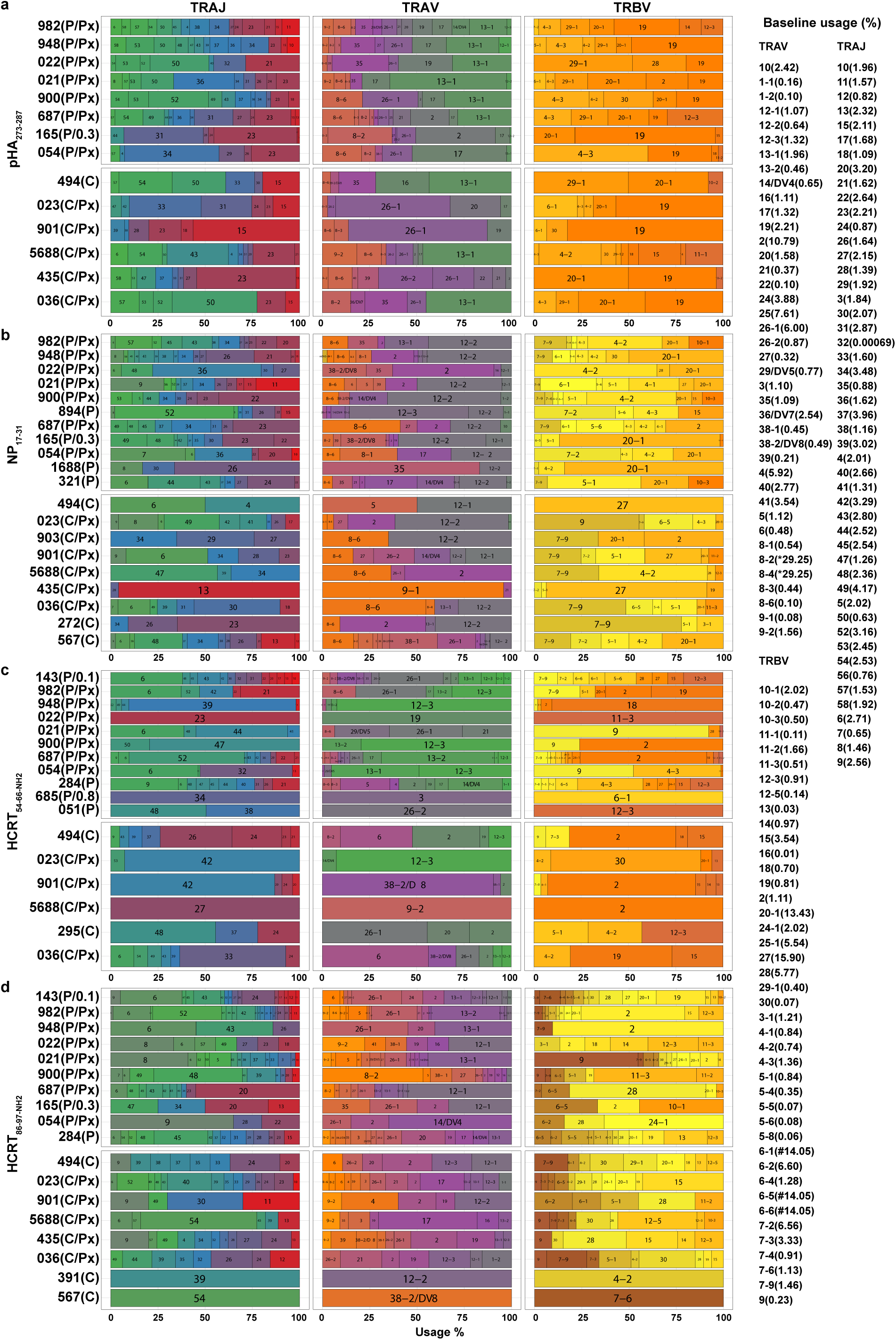
Usage of TCR segment for main pDQ0602 tetramer (pHA_273-287_ (a), NP_17-31_ (b), HCRT_54-66-NH2_ (c) and HCRT_86-97-NH2_ (d)) in patients and controls with corresponding mean usage (%) on blood CD4+ T cells. Note that a few related chains cannot be separated by bulk sequencing but can be distinguished in single cell sequencing; these are denoted by * and # in reference population frequency. P, patient; P/Px, post Pandemrix patient; C, control; C/Px, post Pandemrix control. Interval is shown if patient is early onset (<1.0 year). TCR segment is indicated in box. For subject infomation, see Supplementary Table 9.

**Extended Data Figure 4.**
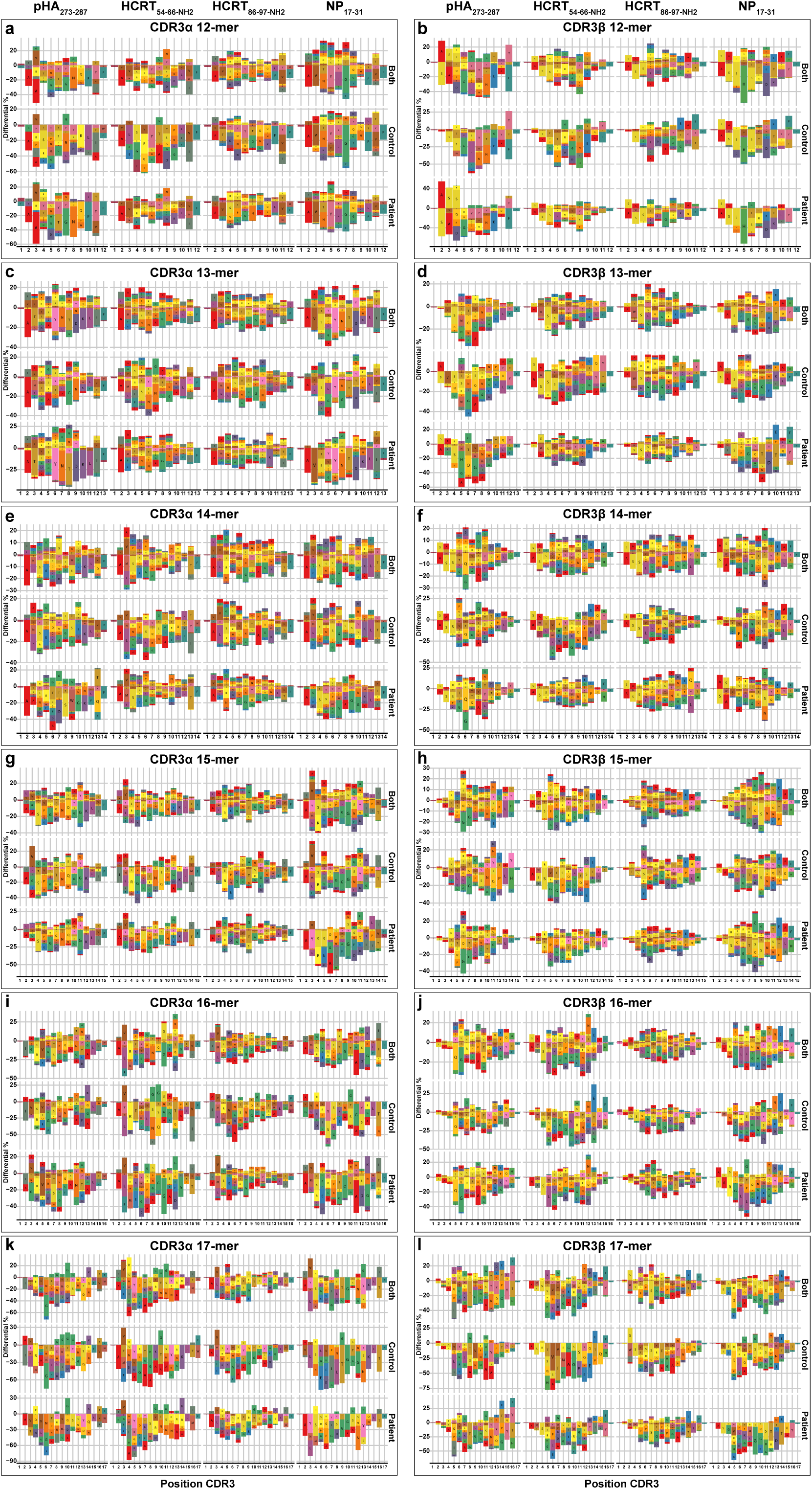
Preferential usage of specific aminoacids in tetramers versus control sequences of various lengths.

**Extended Data Figure 5.**
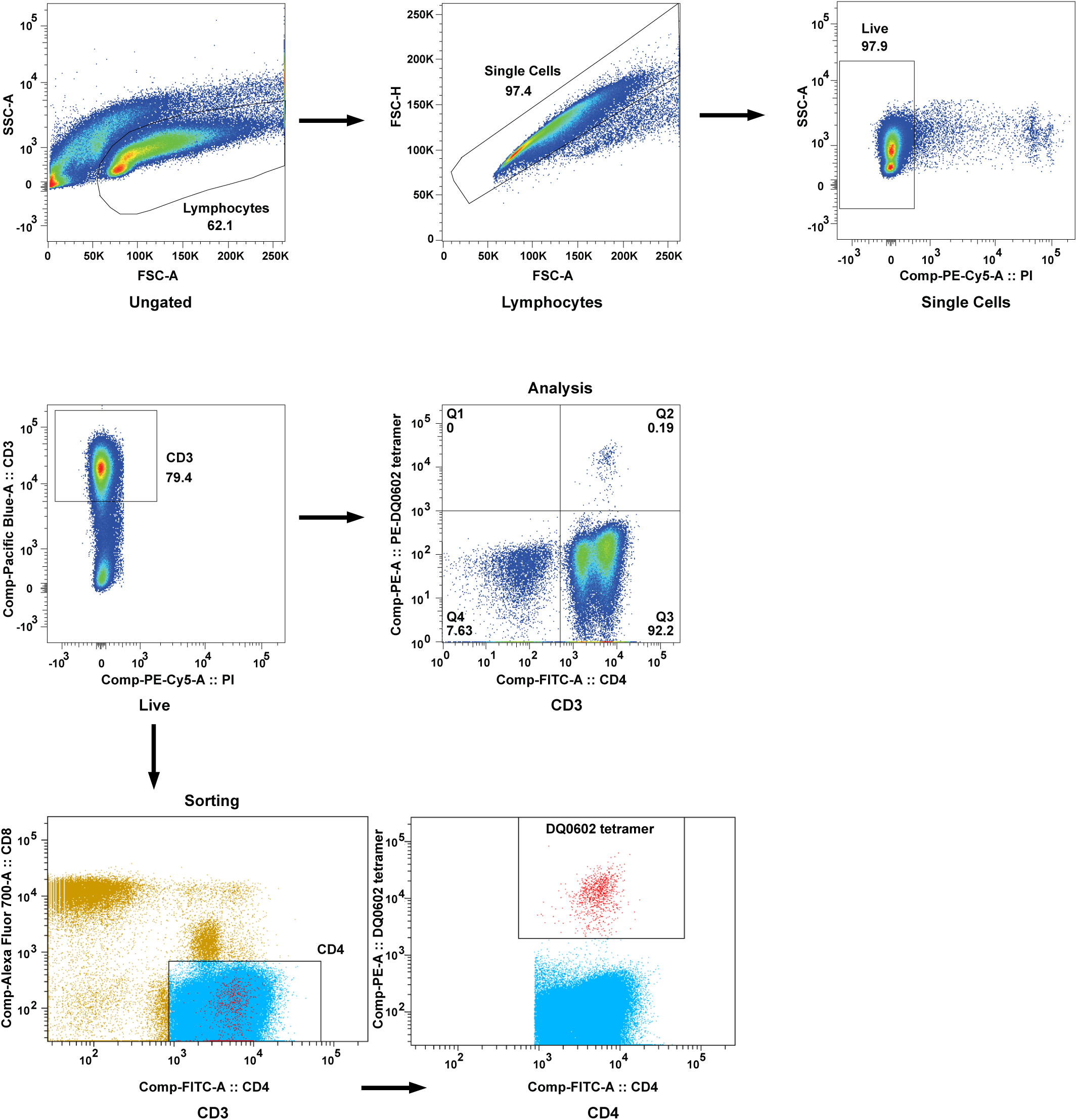
Gate strategy used for analyzing and sorting DQ0602 tetramer positive CD4+ T cells. % parent population is shown. Comp, compensation. A, area. H, height.

